# Repurposing a chromosome segregation ParB-CTPase fold into an ATPase toxin for contact-dependent growth inhibition in plant and animal pathogens

**DOI:** 10.64898/2026.05.05.722872

**Authors:** Jovana Kaljević, Julia E.A. Mundy, Boris Stojilković, Martin Rejzek, Thierry Oms, Frédéric Goormaghtigh, Thomas C. McLean, Ngat T. Tran, Katie E. Johnson, Ananda Sanches Medeiros, Victor Sourjik, Sandra Eltschkner, Laurence van Melderen, Antoine Hocher, Tung B. K. Le

## Abstract

Bacterial competition drives the evolution of antibacterial mechanisms, yet how new activities arise remains poorly understood. A major route to innovation is the reuse of pre-existing genetic systems, whereby conserved protein modules are repurposed in new biological contexts to generate new capabilities. Here, we show that the ParB-CTPase fold, a conserved nucleotide-binding module best known for its role in chromosome segregation, can be functionally repurposed as an antibacterial toxin. We identify ToxB, a ParB-like domain embedded within the polymorphic toxin region of contact-dependent inhibition systems and show that it functions as a potent antibacterial effector. Structural and biochemical analyses reveal that ToxB retains the core architecture of the ParB-CTPase fold but lacks DNA-binding capability and preferentially binds ATP. This shift in nucleotide specificity underpins a distinct mode of action, in which ATP binding and hydrolysis trigger rapid nucleoid compaction, chromosome segregation defects, oxidative stress, cell chaining, and ultimately cell lysis. ToxB also exhibits toxic activity in plant cells, suggesting that it targets conserved cellular processes. Together, these findings provide direct experimental evidence that the ParB-NTPase fold is biologically versatile and can be repurposed for biological roles fundamentally distinct from its ancestral function in DNA segregation.

## Introduction

Bacteria frequently engage in interbacterial warfare to compete for limited resources, yet the evolutionary routes by which new antibacterial activities arise remain poorly understood. A major driver of biological innovation is the reuse of pre-existing genetic systems for new functions^1,2^. Rather than evolving entirely new protein architectures, bacteria frequently repurposed conserved protein modules in different biological contexts, generating new capabilities from ancestral cellular machinery. In several cases, conserved enzymatic activities have been adapted towards interbacterial antagonism. For example, cytidine deaminases, which normally function in the pyrimidine salvage pathway, can evolve into toxins such as DddA that attack genomic DNA and induce mutagenesis in competing cells^3–6^. Similarly, colicins employ a conserved nuclease activity to degrade nucleic acids and disrupt essential cellular processes in competitors^7–9^. These examples highlight the functional plasticity of conserved protein modules and raise the question of how broadly such repurposing occurs across bacterial systems.

One such conserved module is the ParB-CTPase fold, best known for its role in chromosome and plasmid segregation, but increasingly suggested to support diverse biological functions^10–15^. Direct experimental evidence, however, has remained limited^14^. In canonical ParAB*S* systems, ParB binds *parS* DNA sites and, through CTP-dependent conformational switching, assembles sliding clamp-like nucleoprotein complexes that interact with the ATPase ParA to facilitate DNA segregation^10,16–25^. The N-terminal domain of ParB harbors the ParB-CTPase fold, which contains conserved nucleotide-binding motifs, including Box I (C motif) and Boxes II-III (P motifs), that bind the cytosine base and phosphate groups, respectively, enabling CTP binding and hydrolysis^17–20,26,27^. In canonical ParB, this fold is typically coupled to a helix-turn-helix (HTH) domain that together mediates *parS* DNA recognition and higher-order nucleoprotein complex assembly (**Figure 1A**)^10,12,28^. Several ParB homologs, for example, KorB^29^, BisD^30^, VirB^31–33^, and Noc^34^, retain this architecture and act as CTP-dependent molecular switches regulating plasmid segregation, gene expression, or cell division. In contrast, more divergent proteins, such as the eukaryotic sulfiredoxin Srx and the archaeal serine kinase SerK, retain the core ParB-CTPase fold but lack DNA-binding and dimerization domains^35,36^. These proteins do not adopt clamp-like conformations and instead bind ATP to support enzymatic activities unrelated to chromosome segregation. These examples suggest that the ParB-CTPase fold can function as a versatile nucleotide-binding module that supports fundamentally different biochemical activities. Consistent with this idea, our recent bioinformatic survey revealed a far greater diversity of proteins containing the ParB-CTPase fold than previously appreciated^14^. These ParB-like proteins occur in varied domain architectures and genomic contexts, often lacking canonical segregation partners (e.g., ParA), and display substantial variation in nucleotide-binding motifs. In several cases, experimental characterization has shown altered nucleotide specificity, including binding to ATP or GTP rather than CTP^14^. Despite this diversification, their biochemical activities and physiological roles remain largely unexplored.

**Figure 1.**
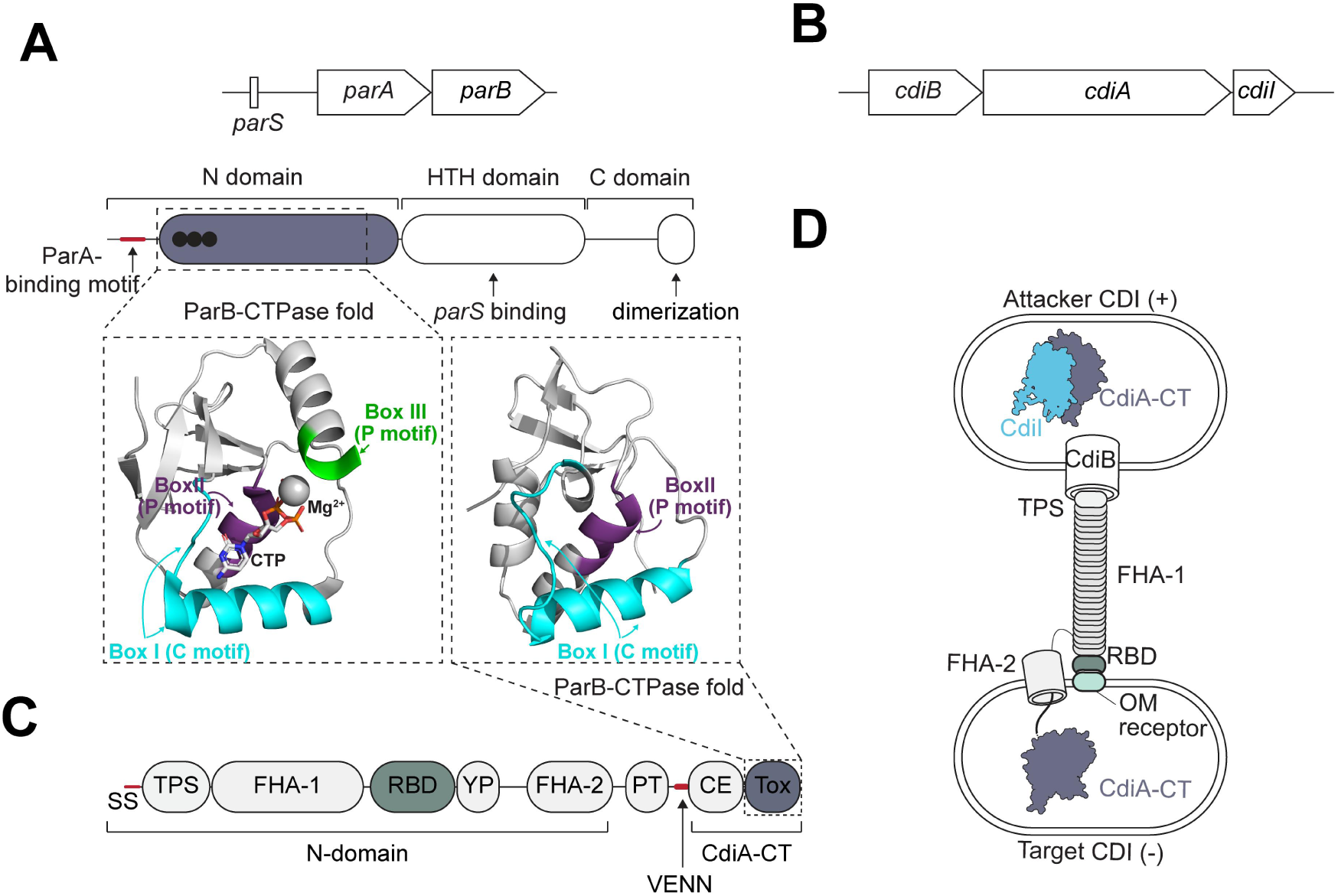
Organization, delivery, and genomic context of ToxB-containing CDI systems. **(A)** Top: Domain organization of canonical ParB proteins. ParB consists of an N-terminal domain containing the ParA-binding motif (small red box) and the ParB-CTPase fold (dashed box; bound CTP shown as black circles), a central helix–turn–helix (HTH) domain mediating *parS* recognition, and a C-terminal dimerization domain. Bottom: structural model of the ParB-CTPase fold (PDB: 7BM8) bound to CTP and Mg²⁺, with conserved nucleotide-binding motifs (Box I, Box II, and Box III) highlighted in cyan, purple, and green, respectively. **(B)** Genetic organization of a canonical contact-dependent inhibition (CDI) locus, consisting of *cdiB*, *cdiA*, and the downstream immunity gene *cdiI*. **(C)** Bottom: domain architecture of the CdiA effector protein. Following the N-terminal signal sequence (SS) and type V secretion system (TPS) domain, CdiA contains FHA-1 repeats, a receptor-binding domain (RBD), YP repeat regions, FHA-2 repeats, and a pre-toxin (PT) module. The variable C-terminal toxin domain (CdiA-CT) is separated by a VENN motif and consists of a short C-terminal entry (CE) region followed by the toxin domain (Tox), which in this case contains ToxB (purple). Top: AlphaFold model of ToxB highlighting the conserved nucleotide-binding architecture, including Box I (C motif, cyan) and Box II (P motif, purple). **(D)** Schematic of CDI-mediated toxin delivery. In CDI⁺ cells, CdiA is secreted through CdiB and displayed on the cell surface. Upon direct cell-cell contact, the RBD engages an outer membrane receptor on the target cell, enabling transfer of the CdiA-CT into the susceptible CDI⁻ cell. The producing cell is protected by the cognate immunity protein CdiI.

To investigate the functional potential of ParB-CTPase fold, we examined a rapidly evolving, previously uncharacterized clade of ParB-like proteins^14^. Within this clade, we identified a ParB-CTPase fold embedded within the C-terminal domain of a CdiA protein. CdiA proteins are toxins of contact-dependent inhibition (CDI) systems, a widespread mechanism used by Gram-negative bacteria to inhibit neighboring cells via direct cell-cell contact^37–40^. CDI systems are composed of three components: CdiB, an outer-membrane transporter; CdiA, a large, secreted protein responsible for toxin delivery; and CdiI, an immunity protein that protects the toxin-producing attacker cell (**Figure 1B**)^41–49^. CdiA has a modular architecture, with a conserved N-terminal region for secretion (SS, TPS, FHA-1 domains) and target-cell recognition (RBD, FHA-2 domains), and a polymorphic C-terminal toxin domain (CdiA-CT), typically separated by a VENN module, that mediates growth inhibition upon delivery into neighboring competitor cells (**Figure 1C-D**)^48,50–54^.

Here, our sequence and structural analysis reveal a clade of CdiA (henceforth, named CdiA-ToxB) that contains a ParB-CTPase fold at its toxin C-terminal domain. However, unlike canonical ParB proteins, which bind CTP and function as DNA-associated molecular switches, CdiA-ToxB lacks DNA-binding activity and binds and hydrolyzes ATP. When delivered into competitor cells as a cleaved toxin domain, CdiA-ToxB acts as a potent growth inhibitor, inducing rapid nucleoid compaction, chromosome segregation defects, and eventually cell death. We suggest that CdiA-ToxB targets conserved cellular processes across kingdoms as ToxB is also toxic to plant cells. Altogether, these findings define a previously unrecognized class of toxins that repurpose the ParB-CTPase fold to disrupt essential cellular functions. More broadly, our results provide direct experimental evidence that diversification of ParB-like fold can generate new biological activities, expanding the role of this conserved fold from genome maintenance to interbacterial antagonism.

## Results

### A divergent ParB-CTPase fold is located at the C-terminal domain of rare bacterial CdiA toxins

Previously, we used a domain-centric search based on the ParB N-terminal domain (PFAM: PF02195) to explore the diversity of ParB-CTPase fold-containing proteins beyond canonical DNA segregation systems^14^. This analysis revealed two major groups. The first comprises a conserved clade of ParB homologs with short branch lengths and canonical domain organization, including a ParA-binding motif, an N-terminal CTP-binding domain, a central helix-turn-helix DNA-binding domain, and a C-terminal dimerization domain (**Figure 2A**, blue and orange dots in the inner and outer rings; example 1). The second is a more divergent clade, characterized by longer branches, with higher sequence variability within nucleotide-binding motifs, and frequent loss of canonical features such as the ParA-binding motif (**Figure 2A**, no orange dots in the outer ring). These features suggest functional diversification beyond chromosome segregation. Within this rapidly evolving group, we identified a distinct subset in which the ParB-CTPase fold is located at the C-terminus of large, modular proteins resembling CdiA toxins (**Figure 2A**, magenta branches, example 2). We termed these proteins CdiA-ToxB and refer to their C-terminal ParB-CTPase fold as ToxB. Analysis of sequence conservation across the ParB-CTPase fold showed that, while canonical ParB proteins retain strong conservation across the Box I (C motif) and Box II-III (P motifs) required for CTP binding, the ToxB group displays altered conservation patterns in these motifs (**Figure 1A, D, Figure 2B, Figure S1A**), suggesting potential changes in nucleotide-binding properties and function.

**Figure 2.**
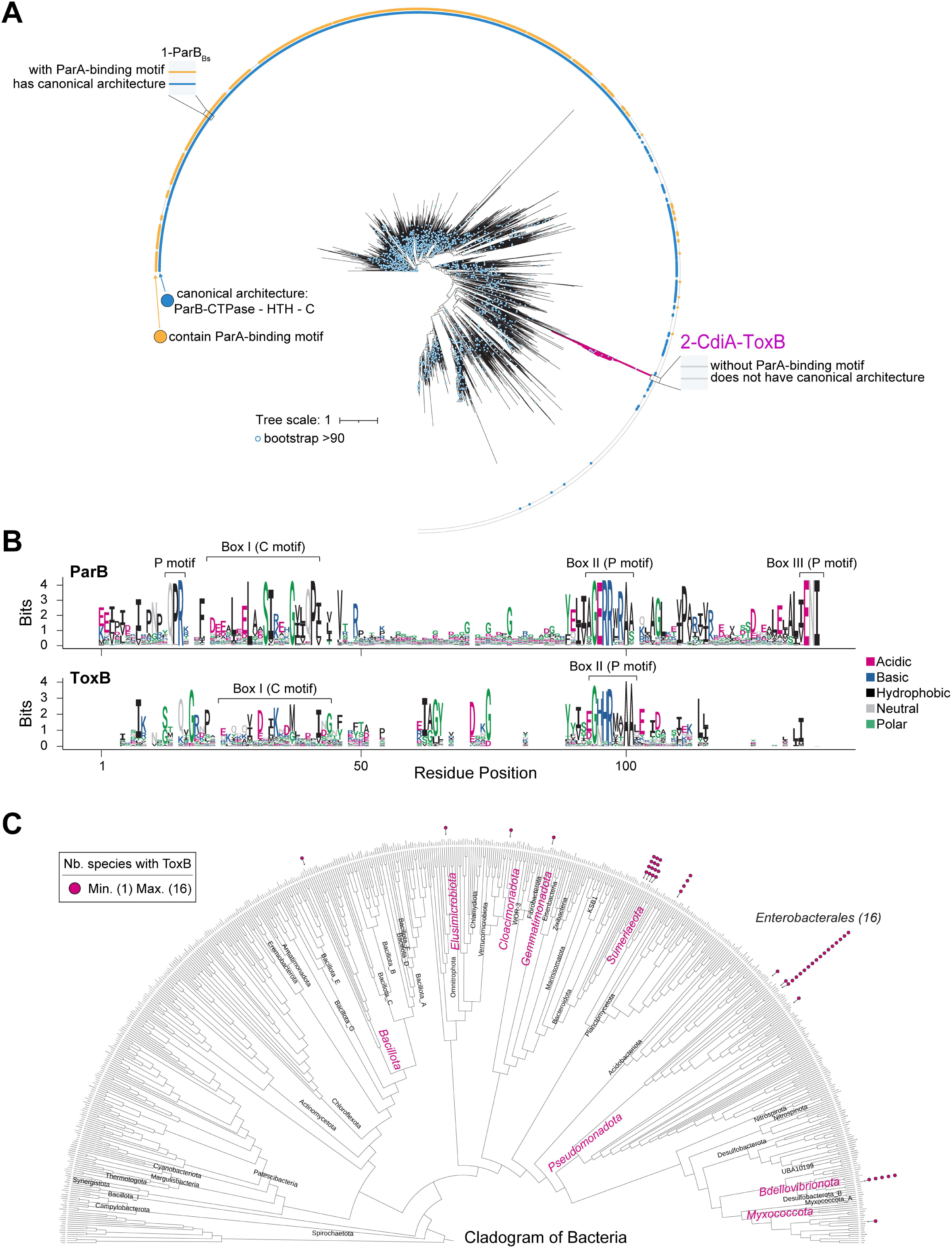
Phylogenetic diversity of ParB-CTPase fold-containing proteins, and ToxB distribution across bacteria. **(A)** Protein domain-level, maximum likelihood phylogenetic tree of 9509 ParB-like domains from bacteria (midpoint rooted); bootstrap support ≥ 90 is indicated by blue dots. Concentric rings summarize key features (from inside to outside): the blue ring indicates canonical domain organization (defined as a positive hit against TIGR00180); the orange ring indicates the presence of the ParA-binding motif (defined as (L|F|I)G[K|R]G(L|F|I). Canonical ParB from *Bacillus subtilis* is shown (example 1). The clade containing CdiA-ToxB proteins is highlighted in magenta (example 2) and corresponds to a distinct group of non-canonical ParB-like domains lacking both the ParA-binding motif and canonical domain organization. **(B)** Sequence conservation and motif architecture of the ParB-CTPase fold. Sequence logos derived from multiple sequence alignments are shown for canonical ParB proteins (top) and ToxB-containing proteins (bottom). Conserved nucleotide-binding motifs are indicated: Box I (C motif), Box II (P motif), and Box III (P motif). **(C)** Phylogenetic distribution of ToxB-containing CDI systems across bacteria. The cladogram represents bacterial species surveyed for the presence of ToxB homologs, summarized at the order level. Magenta dots indicate the number of species encoding ToxB within each order (minimum = 1; maximum = 16). The order with the highest representation of ToxB-containing species, *Enterobacteriales*, is indicated. See also Figure S1.

Next, we examined the phylogenetic distribution and genomic context of CdiA-ToxB homologs (**Figure 2C, Figure S1A**). We found that CdiA-ToxB proteins are rare (identified in only 52 out of 68,709 surveyed bacterial species) but are concentrated within the order Enterobacteriales (16 species), with only sporadic occurrences elsewhere, indicating a restricted yet non-random distribution (**Figure 2C, Figure S1A**). We focused our analysis on four representative Enterobacteriales species of agricultural and clinical relevance: *Erwinia amylovora*, *Providencia rettgeri*, *Klebsiella grimontii*, and *Proteus mirabilis*. In all cases, ToxB is located at the C-terminus of those CdiA proteins, immediately downstream of a VENN or VENN-like motif that typically demarcates the toxic “warhead” region (**Figure S1B**), consistent with canonical CDI organization (**Figure 1C**)^48^. Genomic analysis further showed that these loci reside within conserved *cdiBAI* operons, with *cdiA-toxB* followed by a small gene typically encoding a predicted immunity protein (CdiI, hereafter referred to as iToxB) (**Figure S1A, C**). Notably, ToxB is not uniformly present across strains. While nearly all *E. amylovora* isolates encode *cdiA*-*toxB*, only subsets of *K. grimontii, P. rettgeri,* and *P. mirabilis* isolates carry this locus (**Figure S1D**), consistent with the strain-level variability typical of CDI systems^41,55^. Furthermore, *cdi-toxB*-*itoxB* are occasionally found outside the complete *cdiBAI* operon as truncated or standalone units (**Figure S1A**), a configuration commonly associated with horizontal transfer of CDI toxin-immunity pairs^44,56^. Overall, ToxB contains a divergent ParB-CTPase fold that defines a rare and previously unrecognized class of orphan ParB-like proteins that likely function as CDI toxins.

### ToxB is a potent toxin that inhibits bacterial growth and is neutralized by the immunity protein iToxB

To test whether ToxB inhibits bacterial growth under contact-dependent delivery conditions, we engineered a chimeric CDI system based on the well-characterized *E. coli* EC93 *cdiBAI* locus^50^. We grafted the *toxB-itoxB* toxin-immunity gene pair from *E. amylovora* onto the well-characterized *E. coli* EC93 CDI system under the control of its native promoter. Briefly, the chimeric CDI system was encoded on a plasmid and contained three distinct modules: (i) the EC93 CdiB; (ii) the EC93 CdiA N-terminus fused to the translocation domain of *E. coli* EC869 (YciB), which is known to support the delivery of heterologous CDI toxins into *E. coli* cells, and further fused to ToxB; and (iii) the downstream-encoded iToxB immunity protein. (**Figure 3A**)^43,46^. Thus, all secretion and translocation components were derived from the native EC93-EC869 CDI systems, with only the toxin and immunity modules substituted. In competition assays, the strain carrying the chimeric toxin module resulted in a 10^5^-fold reduction in the viability of susceptible CDI (-) target cells relative to the control without an attacker (**Figure 3A**, top vs. bottom rows). We introduced a point mutation in the conserved arginine residue of the P motif (Box II; R108A), which is typically required for nucleotide binding in ParB-like proteins (**Figure 2B**)^14,19,27^. In contrast to the wild-type toxin, delivery of the R108A variant did not affect target cell viability (**Figure 3A**, middle vs. bottom rows), suggesting that an intact nucleotide-binding site is essential for ToxB toxicity. These findings established that CdiA-ToxB functions as a *bona fide* CDI toxin whose activity depends on its ParB-like nucleotide-binding fold.

**Figure 3.**
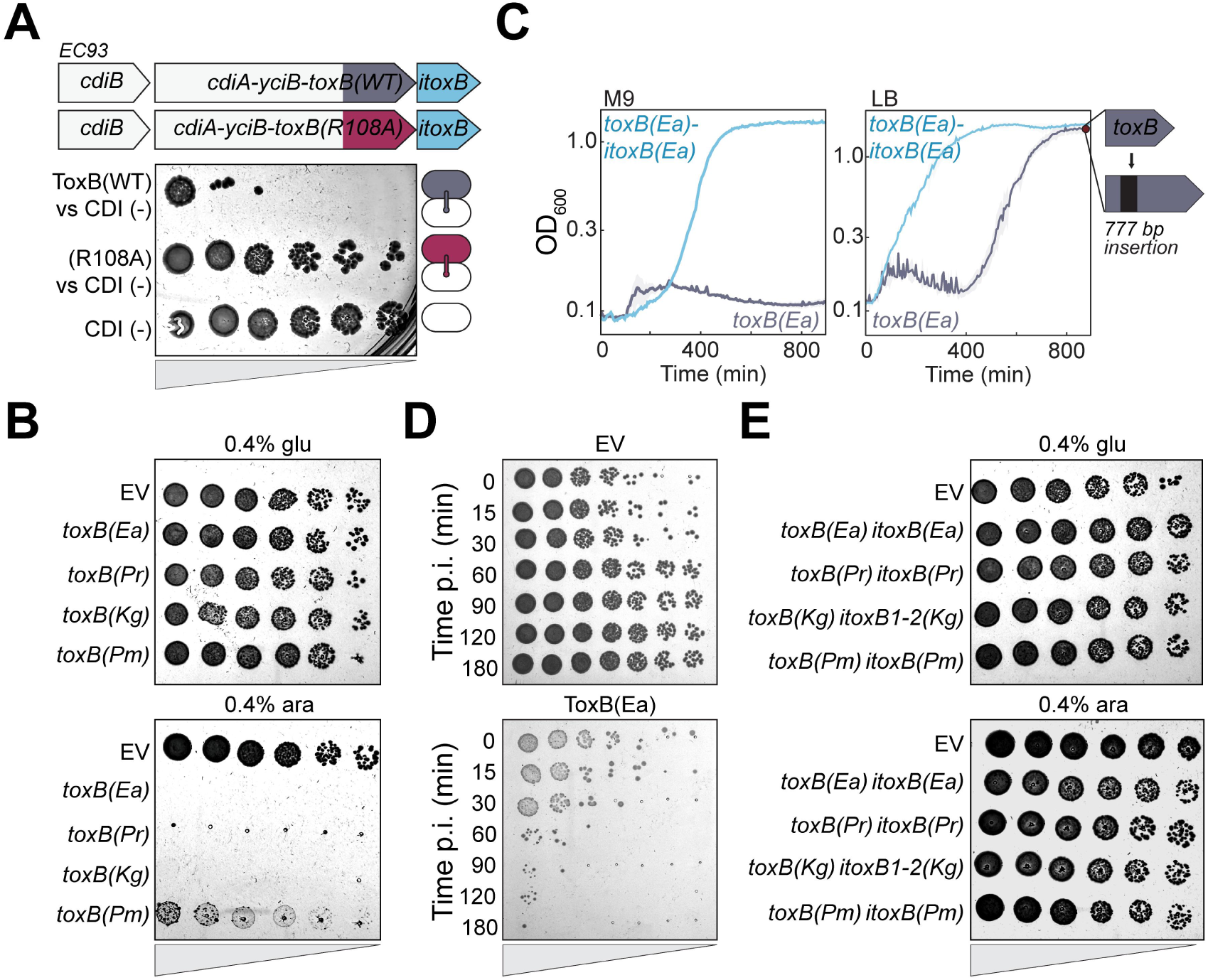
ToxB acts as a bacterial toxin, and iToxB as its immunity partner. **(A)** CDI-mediated delivery of the ToxB and ToxB ParB-CTPase fold mutant. Top: schematic of the chimeric two-component *cdiA-cdiB* contact-dependent inhibition (CDI) system derived from *E. coli* EC93. The *toxB(WT)* and *toxB(R108A)* coding sequences (residues 7984-8437 of *Erwinia amylovora*; dark purple and magenta, respectively) were fused to the C-terminus of *E. coli* CdiA^EC^^93^-YciB, downstream of the translocation domain. The cognate immunity gene (*itoxB*, light purple) was placed immediately downstream of *cdiA* to protect producing (attacker) cells. Bottom: serial dilution spotting assay of target cells following co-culture with attacker strains. A CDI-negative control shows normal growth of prey cells (bottom row). **(B)** Spot dilution assays of *E. coli* cells expressing ToxB proteins from different species. Cells harboring an empty vector (EV) or expressing *toxB* homologs from *Erwinia amylovora toxB(Ea),* Uniprot: A0ABX7MJ29, *Providencia rettgeri toxB(Pr),* Uniprot: A0A9N8GWP0, *Klebsiella grimontii toxB(Kg),* UniProt: A0ABD7AHS9, or *Proteus mirabilis toxB(Pm)*, UniProt: A0A6I7DA34 from the pBAD33 vector were serially diluted and spotted onto LB agar supplemented with either 0.4% glucose (glu; repressing conditions) or 0.4% arabinose (ara; inducing conditions). **(C)** Growth curves of *E. coli* cells expressing *toxB(Ea)* alone or together with its cognate immunity gene, *itoxB(Ea),* in M9 (left) or LB (right) medium. Survivors repeatedly carried a 777-bp insertion within a *toxB* gene (777 bp insertion). **(D)** Time-course viability assay following induction of *toxB(Ea)* expression. *E. coli* cells carrying an empty vector or pBAD33-*toxB(Ea)* were induced with 0.4% arabinose and sampled at the indicated times post-induction (p.i.). **(E)** Cognate immunity proteins iToxB(Ea), iToxB(Pr), iToxB(Kg1), iToxB(Kg2), iToxB(Pm). UniProt codes: A0ABX7MJ22, A0A9N8CZ08, A0A285B4B9, A0ABD7AI9, and A0A6I7D2M4, respectively) protect against ToxB-mediated toxicity. *E. coli* cells expressing the native context operon *toxB-itoxB* genes were grown under inducing conditions, serially diluted and spotted onto LB agar supplemented with either 0.4% glucose (glu; left), or 0.4% arabinose (ara; right). See also Figure S2.

To examine the intrinsic toxicity of ToxB independently of CDI-mediated delivery, we expressed *toxB* directly in the cytoplasm of *E. coli* from a tightly regulated arabinose-inducible promoter on a low-copy-number plasmid. Production of ToxB from four representative species (*E. amylovora*, *P. rettgeri*, *K. grimontii*, and *P. mirabilis*) caused a complete loss of *E. coli* viability in both solid and liquid media (**Figure 3B-C**). Growth inhibition occurred in both rich (LB) and minimal (M9) media, with sustained inhibition in M9 and late recovery in LB, due to spontaneous insertions disrupting the *toxB* sequence (**Figure 3C**). ToxB toxicity was highly potent, as a brief 15 min induction pulse was sufficient to cause a markedly reduction in viability (**Figure 3D, Figure S2A**). Moreover, induction at arabinose concentrations 1000-fold lower than a standard level still markedly impaired growth (**Figure S2B**). The *P. mirabilis* toxin displayed reduced potency relative to the other variants, but nevertheless still significantly impaired growth compared to empty-vector controls (**Figure 3B; Figure S2A**). In all cases, growth inhibition was fully rescued by co-expression of the downstream gene *itoxB*, both on solid and in liquid media (**Figures 3C, E,** and **S2C**), confirming its role as an immunity protein. This protection was specific, as non-cognate toxin-immunity pairs failed to completely restore growth (**Figure S2D**). Together, these results demonstrate that ToxB is an intrinsically potent toxin whose activity is neutralized by the cognate immunity protein iToxB, and that the ParB-like fold is sufficient to inhibit the growth.

### Despite harboring a ParB-CTPase fold, ToxB binds and hydrolyzes ATP instead

ToxB belongs to a rapidly evolving clade of ParB-CTPase fold-containing proteins, recently recognized as a versatile nucleotide-binding module that binds nucleotides beyond CTP, including ATP and GTP^14^. We therefore asked whether the catalytic core of ToxB remains functional and, if so, which nucleotide it binds. To identify the associated ligand, we purified the *E. amylovora* ToxB-iToxB complex following heterologous expression in *E. coli*, then denatured the proteins to release any bound small molecules and analyzed the extract by LC-MS/MS (**Figure 4A**). Unexpectedly, we detected ATP and ADP, but not CTP, at approximately 1:1 stoichiometry with ToxB (**Figure 4B**), indicating that ToxB preferentially binds ATP.

**Figure 4.**
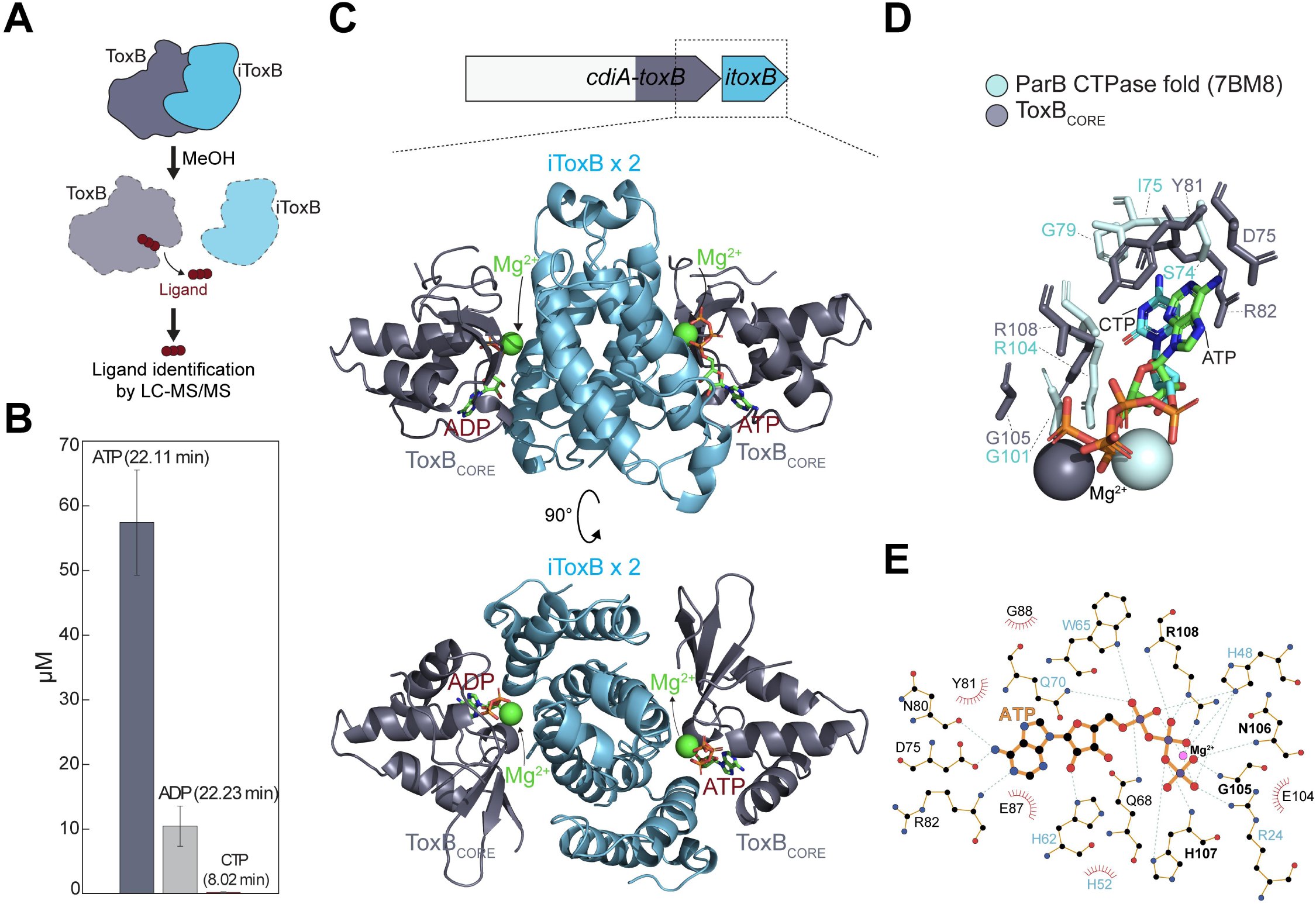
ToxB is a ParB-like protein that binds ATP. **(A)** Schematic of the ligand identification workflow using LC-MS/MS. **(B)** Quantification of nucleotides detected by LC-MS/MS following dissociation of the ToxB-iToxB complex. ATP, ADP, and CTP levels are shown at the indicated retention times. **(C)** Crystal structure of the *E. amylovora* ToxB-iToxB complex, at 2.4 Å resolution. Top, schematic of the *toxB-itoxB* operon; regions cloned for protein purification are highlighted in purple. Top and bottom views of the crystal structure of ToxB_CORE_ (dark purple) in complex with its cognate immunity, iToxB (blue). Two ToxB_CORE_ protomers are shown bound to ATP or ADP, as indicated; Mg²⁺ ions are shown as spheres (green). **(D)** Architecture of the nucleotide-binding pocket. Zoom-in view of the ToxB_CORE_ (purple) nucleotide-binding site showing residues coordinating ATP and Mg²⁺. The pocket is overlaid with the CTP-bound N-terminal ParB-CTPase fold from *C. crescentus* ParB (PDB 7BM8; cyan). Conserved nucleotide-binding residues are indicated. **(E)** LigPlot representation of ATP binding within the ToxB active site. Schematic interaction map showing contacts between ATP and residues from the toxin and the immunity protein (blue). Hydrogen bonds are indicated by green dashed lines and hydrophobic contacts by red arcs. See also Figure S3 and Supplementary Table 1.

To uncover the structural basis of ATP binding, we determined the co-crystal structure of the *E. amylovora* ToxB-iToxB complex to 2.4 Å resolution using a truncated toxin variant comprising only the catalytic core (ToxB_CORE_), lacking the C-terminal cytoplasm entry helix immediately downstream of the VENN sequence (**Figure 4C**). This truncation, while retaining full toxicity, substantially improved protein yield and crystallization. The structure reveals an iToxB dimer bound by two flanking ToxB_CORE_ monomers, one on each side (**Figure 4C**). Clear electron density corresponding to nucleotides was observed in the active site of each ToxB monomer, with one monomer bound to ATP and the other to ADP (**Figure 4C-E**).

Structural comparison with the canonical CTP-bound ParB showed that the overall architecture of the phosphate-binding region is highly conserved (**Figure 4D-E)**^17,18,20^. Residues surrounding the nucleotide phosphates occupy analogous positions, and the Mg²⁺ ion is coordinated in a similar geometry (**Figure 4E-D**). In both proteins, the phosphate groups are bound by a cluster of residues that form the GxxR phosphate-binding motif. In ToxB, residues G105, N106, H107, and R108 occupy positions equivalent to phosphate-binding residues in canonical ParB (G101, E102, R103, and R104 in PDB model 7BM8) (**Figure 4D**, purple ToxB vs cyan ParB amino acids)^18^. Moreover, in both complexes, the base is bound by a network of hydrogen bonds involving residues S74 and the peptide-backbone atoms of G79 (O), L81 (N) and Q82 (N) in ParB-CTP for cytosine binding (**Figure 4D**, residues in cyan)^18^, and in ToxB, residue D75 and the peptide backbone atoms of N80 (O) and R82 (N) for adenine binding (**Figure 4D-E**). There are, however, notable differences in base recognition between the ToxB and canonical ParB (**Figure 4D-E**). Besides the substitution of a serine (S74) in ParB-CTP by an aspartate residue (D75) in ToxB, forming a hydrogen bond with N4 of the cytosine or N6 of the adenine, respectively, the ring plane of the cytosine base in ParB-CTP is located in a mainly hydrophobic pocket. This pocket comprises residues L71, I75, V80, L81, and the hydrophobic part of Q82 of the nucleotide-binding subunit, as well as I134 and the hydrophobic part of Q138 from the respective other monomer within a maximum radius of 4.0 Å^18^. In contrast, the adenine base in ToxB is engaged in multiple 𝜋-stacking interactions: the benzene ring is inserted between the side chain of Y81 (parallel-displaced π-stacking) and R82 (cation-π stacking), while the pyrrole ring T-stacks with W65 from the adjacent iToxB monomer (**Figure 4D-E**). Additional residues from the immunity protein also contribute to nucleotide binding (R24 from iToxB monomer A, and Q70, H48, and H62 from iToxB monomer B); however, they are primarily involved in interactions with sugar-phosphate moieties. An opposite ToxB’s active site containing ADP displays essentially an equivalent network of interactions (**Figure S3A, Supplementary Table 1**).

During size-exclusion chromatography, a fraction of *E. amylovora* ToxB-iToxB complex dissociated, yielding a fraction of ToxB alone that was subsequently used for biochemical characterization in the absence of inhibitory immunity protein. Microscale thermophoresis (MST) confirmed specific ATP binding with a moderate dissociation constant (K_D_) of ∼87 μM, and no detectable binding to other NTPs (**Figure 5A**). As a control, mutation at the conserved arginine in the P-motif (R108A) severely impaired ATP binding (K_D_ ∼522 mM).

**Figure 5.**
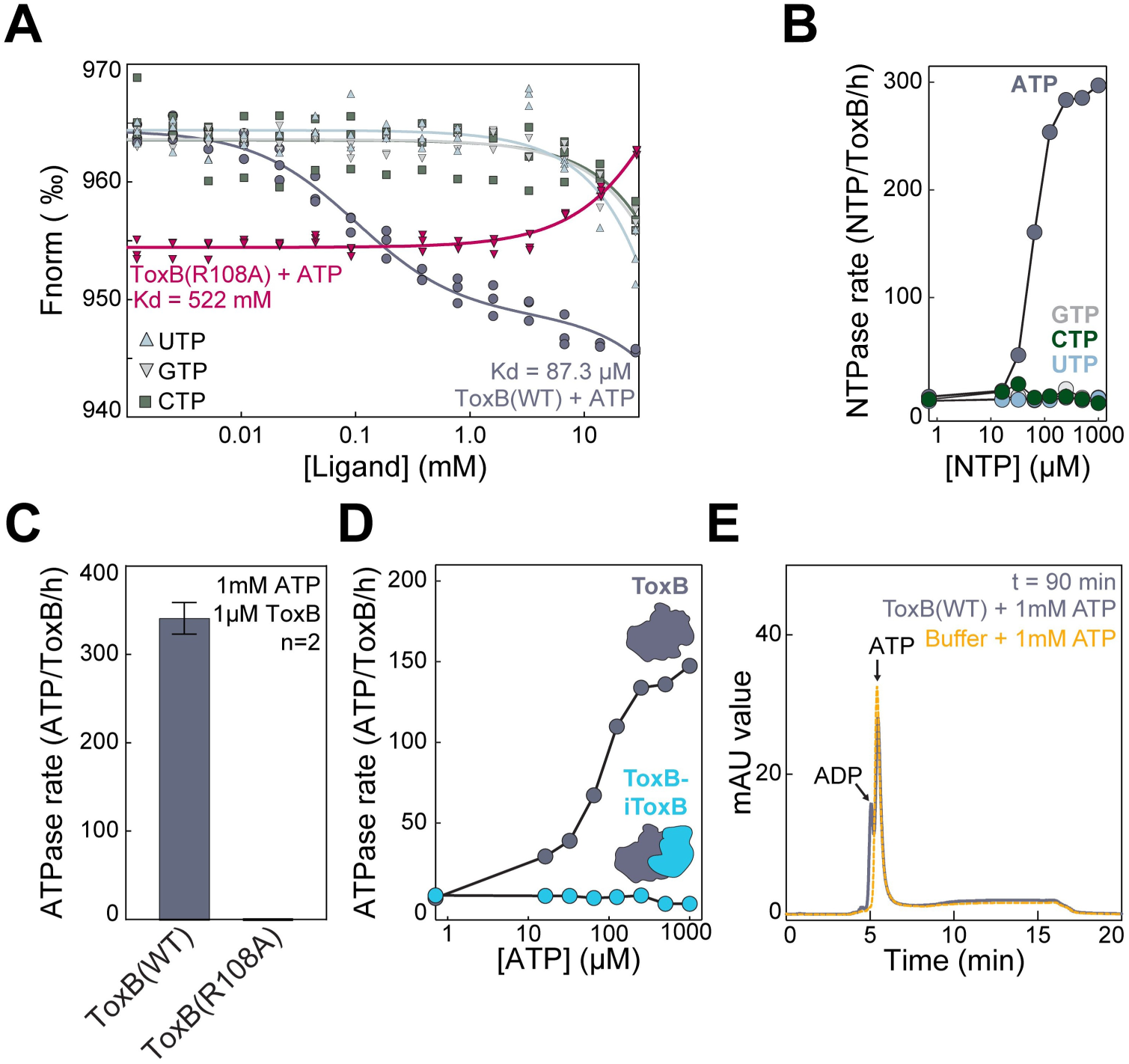
ToxB is a ParB-like ATPase that binds and hydrolyzes ATP. **(A)** ToxB binds ATP, and this interaction is mediated by the conserved ParB-CTPase nucleotide-binding fold. Microscale thermophoresis (MST) analysis of nucleotide binding to ToxB. Normalized fluorescence (Fnorm) is plotted against nucleotide concentration for ToxB(WT) with ATP (dark purple), UTP (blue), GTP (grey), and CTP (green), and for the ToxB(R108A) mutant with ATP (magenta). Apparent equilibrium dissociation constants (Kd) are indicated. **(B)** ToxB preferentially hydrolyzes ATP. NTP hydrolysis rates were measured at increasing concentrations (1-1,000 µM) of ATP, CTP, GTP, or UTP. **(C)** The GxxR motif is required for ATP hydrolysis. ATP hydrolysis rates of ToxB(WT) and ToxB(R108A) were measured at 1 mM ATP. **(D)** The immunity protein inhibits ATP hydrolysis. ATP hydrolysis rates for ToxB were measured at increasing ATP concentrations (1-1,000 µM) in the presence or absence of the cognate immunity protein iToxB. **(E)** ATP is converted to ADP *in vitro*. Representative SAX-UV chromatograms showing nucleotide composition after 90 min incubation of ToxB with ATP. ATP and ADP peaks are indicated. Buffer-only controls supplemented with ATP are shown in red. See also Figure S3.

Moreover, the presence of ADP in one ToxB monomer suggested that ToxB hydrolyzes ATP. Indeed, ToxB protein alone hydrolyzed ATP *in vitro*, with no detectable activity towards other NTPs (**Figure 5B)**. Under saturating ATP concentrations, ToxB hydrolyzed ∼300 ATP molecules per enzyme per hour, corresponding to a slow but sustained ATP turnover (**Figure 5B**). In contrast, the ToxB(R108A) variant, unable to bind ATP, did not detectably hydrolyze ATP (**Figure 5C**). ATP hydrolysis was abolished in the ToxB-iToxB complex (**Figure 5D**), consistent with structural occlusion of the active site by the immunity protein (**Figure 4C, E, Figure S3A**).

Because several well-known ATP-dependent toxins synthesize modified nucleotide derivatives rather than simply hydrolyzing ATP^57,58^, we further analyzed ToxB reaction products using anion chromatography with SAX-UV detection. Only ATP-to-ADP conversion was detected, with no formation of adenosine oligophosphate (tetra-/penta-) nucleotides even after prolonged incubation (**Figure 5E, Figure S3B**), suggesting that ToxB functions as an ATPase, at least in the presence of ATP alone. Altogether, these findings demonstrate that ToxB repurposes a conserved ParB-CTPase fold to function as an ATP-dependent toxin whose catalytic activity is inhibited by its immunity protein through direct protein-protein interaction.

### ToxB intoxication causes rapid nucleoid collapse prior to membrane permeabilization

To determine how ToxB ATPase activity affects cells *in vivo*, we monitored *E. coli* expressing *toxB(WT)* at the single-cell level using a microfluidics device that enabled precise control of toxin induction. Shortly after arabinose addition, cells displayed moderate elongation and chaining, eventually culminating in cell rupture (**Figure 6A**, white arrowheads**; Movie S1**). Loss of a constitutively expressed cytosolic mCherry signal confirmed release of intracellular contents upon lysis (**Figure S4A, Movie S2**). Consistently, a membrane-impermeable dye, propidium iodide (PI), accumulated in ToxB-producing cells but not in control cells (**Figure S4B**), further indicating loss of membrane integrity at late stages. However, we observed that both ToxB and its immunity protein localized exclusively to the cytoplasm when produced as fluorescent fusions (**Figure 6B, Figure S4C**). As neither protein contains predicted membrane-targeting features and both are cytosolic, cell lysis is unlikely to be a direct effect of the toxin but instead a downstream consequence of intracellular damage.

**Figure 6.**
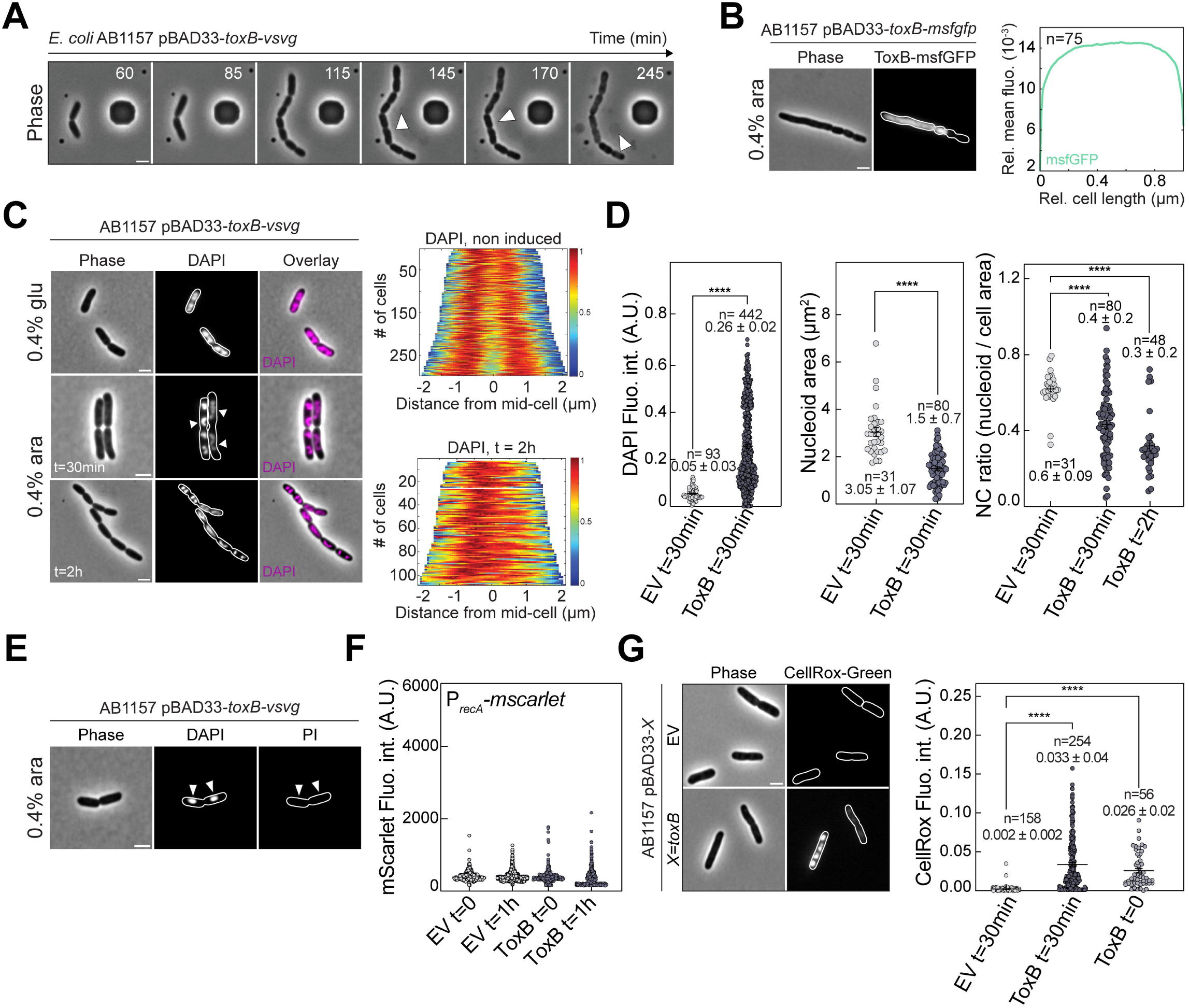
ToxB intoxication induces rapid chromosome compaction and oxidative stress prior to membrane permeabilization. **(A)** Phase-contrast microscopy of *E. coli* AB1157 cells expressing *toxB(WT)* from pBAD33-*toxB-vsvg* following induction in a microfluidics flow cell. Images were acquired every 5 min. Arrowheads indicate progressive morphological changes and release of cytosolic content. See also Movie S1. **(B)** ToxB localizes to the cytosol. Representative phase-contrast and fluorescence images of AB1157 cells expressing *toxB-msfgfp* from pBAD33. Right, mean pole-to-pole fluorescence intensity profiles of msfGFP signal. **(C)** ToxB induces nucleoid compaction. Left: representative phase-contrast, DAPI fluorescence, and overlay images of AB1157 cells expressing *toxB* from pBAD33-*toxB-vsvg* under non-induced conditions and following *toxB* induction at the indicated times. Right: demographs of DAPI fluorescence intensity along the long cell axis, centered at mid-cell, for non-induced cells (top) and cells 2 h after induction (bottom). Heatmaps represent relative fluorescence intensities. **(D)** Quantification of nucleoid perturbation. Left: single-cell DAPI fluorescence intensity measured 30 min after induction compared to empty vector (EV) controls; images were acquired under identical settings. Middle: nucleoid area per cell. Right: ratio of nucleoid area to total cell area (NC ratio) measured at 30 min and 2 h post-induction. Mean ± SD and number of cells analyzed (n) are indicated. Statistical significance was assessed using unpaired two-tailed Student’s *t*-tests or one-way ANOVA with multiple-comparisons testing, as indicated (****, *P* < 0.0001). **(E)** Membrane permeabilization occurs at later stages of intoxication. Representative phase-contrast, DAPI, and propidium iodide (PI) fluorescence images of AB1157 *E. coli* cells expressing *toxB* from pBAD33-*toxB-vsvg.* The PI signal indicates loss of membrane integrity at later time points, when DNA compaction is already apparent (arrowheads). **(F)** ToxB does not activate the SOS response. Single-cell fluorescence intensity of the SOS reporter P*_recA_-mScarlet* in MG1655 *E. coli* cells expressing *toxB* from pBAD33-*toxB-vsvg* or empty vector at the indicated times after induction. **(G)** ToxB activates intracellular oxidative stress. Left: representative phase-contrast and CellROX-Green fluorescence images of AB1157 *E. coli* cells expressing *toxB* from pBAD33-*toxB-vsvg,* or empty vector. Right: quantification of CellROX-Green fluorescence intensity under non-induced, induced, and repressed conditions. Mean ± SD and n values are indicated. Unless otherwise indicated, experiments were performed using *Erwinia amylovora* ToxB(Ea). For all, induction and repression were performed with 0.4% arabinose and 0.4% glucose, respectively. For all, cell outlines were generated using Oufti; scale bars, 2 µm. See also Figure S4.

Cell elongation and chaining suggested perturbed cell-cycle progression and chromosome segregation^59–61^. Indeed, using a chromosomally encoded HupA-GFP fusion, we observed that chromosome segregation rapidly failed following ToxB induction. Instead of resolving into two distinct nucleoids that move toward opposite halves of the cell, chromosomes progressively collapsed into a single compact mass that persisted as cells continued to elongate (**Figure S4D, Movie S3**). This condensed state remained stable throughout the time course, with HupA-GFP signal declining only upon membrane damage and cell lysis. Consistent with live-cell imaging, DAPI staining revealed a rapid nucleoid collapse within 30 minutes of induction (**Figure 6C**), followed by the appearance of aberrant structures, including threads, loops, and dense foci at later time points (**Figure 6C**, 2 h). Quantitative analysis showed a significant increase in DAPI signal intensity alongside reduced nucleoid area and nucleoid-to-cell ratio, indicative of chromosome condensation (**Figure 6D**). Importantly, nucleoid condensation consistently preceded PI uptake across all four ToxB homologs tested (**Figure 6E, Figure S4E**), indicating that defects in chromosome organization and segregation occur prior to membrane permeabilization.

Despite severe chromosomal defects, ToxB expression did not trigger canonical DNA damage responses^61–64^. And unlike several previously described CdiA toxins, ToxB did not behave as a nuclease^8,44,52,54,65^; genomic DNA remained uncleaved for at least two hours following toxin induction (**Figure S4F**), and DAPI staining still indicated the presence of double-stranded DNA within lysed cells after five hours (**Figure S4G**). Consistent with the lack of DNA degradation, an SOS reporter (P*_recA_-mScarlet*) in which the DNA damage-inducible *recA* promoter drives *mScarlet* expression showed no increase in fluorescence upon ToxB expression (**Figure 6F**). In contrast, the CellROX Green reporter showed a strong increase in intracellular reactive oxygen species following ToxB induction (**Figure 6G**). Notably, an elevated CellROX level was detectable as early as 15 minutes post-induction, before any visible morphological changes such as cell elongation or chaining. Oxidative stress is, therefore, also an early event in ToxB intoxication. Altogether, these results show that ToxB disrupts chromosome organization and segregation, induces early oxidative stress, and ultimately leads to membrane failure and lysis.

### The ATP-binding pocket is essential for ToxB toxicity

We next tested whether ATP binding by the ParB-like fold is required for ToxB toxicity. Potential functionally important residues were selected by integrating data from both sequence conservation (**Figure S5A**) and the ToxB-ATP-iToxB structure (**Figure S5B**) and were subsequently substituted by alanine. All alanine-substituted variants were expressed at levels comparable to wild-type ToxB (**Figure S5C**), permitting direct functional comparison.

Growth assays in both solid and liquid media grouped mutants into three classes (**Figure 7A; Figure S5D**). Class I mutants retained wild-type toxicity (**Figure 7A**, grey and **Figure S5D,** circle), Class II mutants showed partial attenuation (**Figure 7A**, purple and **Figure S5D**, triangle), and Class III mutants completely abolished toxicity (**Figure 7A**, magenta, **Figure 7B, Figure S5D**, arrow). Class III mutants included substitutions in residues that directly contact ATP, such as those in the C-motif (M76, Y81) and the P motif, GxxR (G105, R108), as well as a ToxB-specific residue (W134) (**Figure 4E, Figure S5A-B**). Importantly, all non-toxic Class III mutants retained interaction with the immunity protein iToxB (**Figure S5E**).

**Figure 7.**
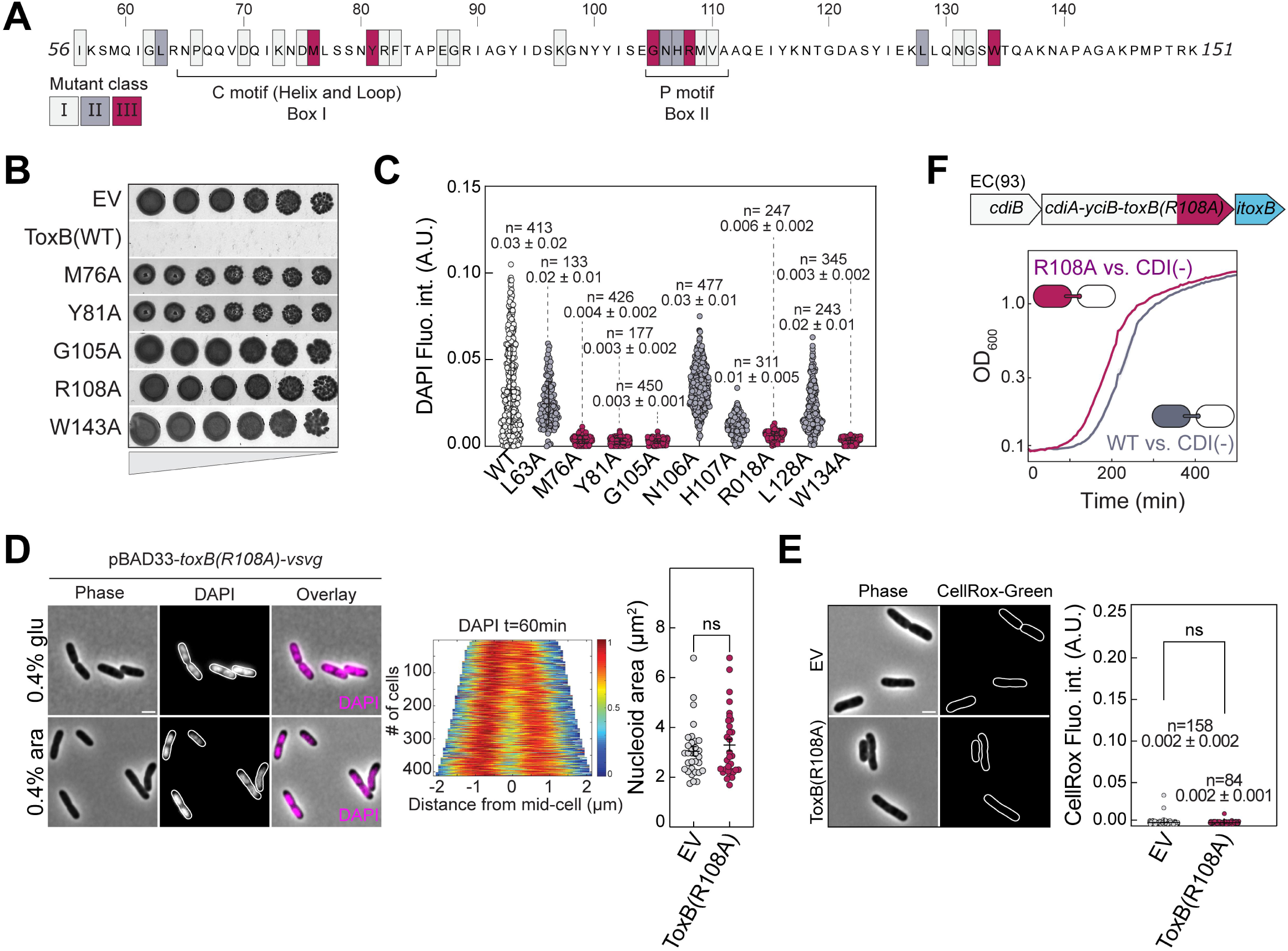
Structure-guided mutational analysis links the ParB-CTPase fold to ToxB toxicity. **(A)** Structure-guided mutagenesis defines functional classes of ToxB mutants. Residues selected for alanine mutagenesis (highlighted) were chosen based on a combination of sequence conservation across ToxB homologs and canonical ParB-CTPase fold from *Caulobacter* ParB (See also Figure S5A) and structural correspondence to the ParB-CTPase fold. Residues are color-coded according to mutational outcome: class I, no loss of toxicity (grey); class II, partial loss of toxicity (purple); class III, complete loss of toxicity (magenta). Toxicity classification was based on growth inhibition and live-cell imaging assays (see panel B and Figures S5-S6). **(B)** P and C motif mutations abolish growth inhibition. Spot dilution assays of AB1157 cells expressing empty vector (EV), ToxB(WT), or representative Class III mutants (M76A, Y81A, G105A, R108A, W134A) under inducing conditions (0.4% arabinose). **(C)** Loss of toxicity correlates with the absence of nucleoid compaction. Single-cell DAPI fluorescence intensity quantified for ToxB(WT), Class II mutants, and Class III mutants. Mean ± SD and number of cells analyzed (n) are indicated. **(D)** The GxxR mutant ATP-binding defective mutant, ToxB(R108A), fails to perturb nucleoid organization. Left, representative phase-contrast, DAPI, and overlay images of AB1157 cells expressing *toxB(R108A)* or empty vector under non-induced and induced conditions. Center, DAPI demographs (right) show no nucleoid disorganization upon induction. Right, quantification of nucleoid area per cell demonstrates no significant difference between R108A-expressing cells and empty vector controls (ns). **(E)** The ToxB(R108A) mutant does not induce oxidative stress. Representative phase-contrast and CellROX-Green fluorescence images (left) and quantification of CellROX-Green fluorescence intensity (right) demonstrate no increase in intracellular oxidative stress compared to empty vector controls. Mean ± SD and n values are indicated. **(F**) The ToxB(R108A) mutant fails to inhibit growth of prey cells in liquid competition assays. Top, schematic of the chimeric two-component *cdiA-cdiB* contact-dependent inhibition (CDI) system derived from *E. coli* EC93, as in Figure 3A. Bottom, liquid competition assays monitoring target cell growth (OD_600_) over time. The cartoon depicts the attacker and prey cells used in the assay. Unless otherwise indicated, experiments were performed using *Erwinia amylovora* ToxB(Ea). See also Figures S5-6.

Single-cell imaging directly linked ATP-binding capacity to cellular outcome. Class I mutants recapitulated wild-type phenotypes, including nucleoid compaction, chromosome segregation defects, and extensive cell chaining (**Figure S6A**). Despite partial recovery in bulk growth, Class II mutants still induced nucleoid condensation and segregation defects (**Figure 7C, Figure S6B**). Time-lapse imaging of the representative Class II mutant N106A showed rapid nucleoid collapse followed by failed chromosome segregation and eventual cell death (**Figure S6D, Movie S4**). In contrast, Class III mutants, including the catalytic mutant ToxB(R108A), displayed a normal chromosome organization, with diffuse, well-segregated nucleoids, similar to the empty vector control, and no evidence of cell chaining (**Figure 7C-D**, and **Figure S6C**). Mutant ToxB(R108A) also failed to induce oxidative stress as measured by CellROX staining (**Figure 7E**). Moreover, when delivered to target cells via the CDI chimera that mimics physiological transfer, ToxB(R108A) exhibits reduced inhibitory activity, with improved target cell growth relative to wild type, even under liquid conditions where cell-cell contact and thus CDI efficiency are limited (**Figure 7F, Figure 3A**). Together, these results demonstrate that ATP binding by the ParB-like fold is essential for ToxB toxicity, and disruption of nucleotide binding eliminated ATPase activity, preventing chromosome collapse and oxidative stress.

### ToxB drives progressive cellular failure and exhibits cross-kingdom toxicity

To determine whether ToxB directly depletes intracellular ATP, we monitored ATP levels using a genetically encoded reporter whose fluorescence signal reflects intracellular ATP levels^66^. ATP levels remained largely stable during the early stages of ToxB production and declined only at later timepoints, when the cells lost membrane integrity (∼1 h after induction) (**Figure 8A-B**). This delayed ATP depletion contrasts with the rapid onset of cellular defects such as nucleoid condensation and oxidative stress, indicating that ATP depletion is a downstream consequence of ToxB’s mode of action rather than an initiating event, consistent with the slow rate measured earlier.

**Figure 8.**
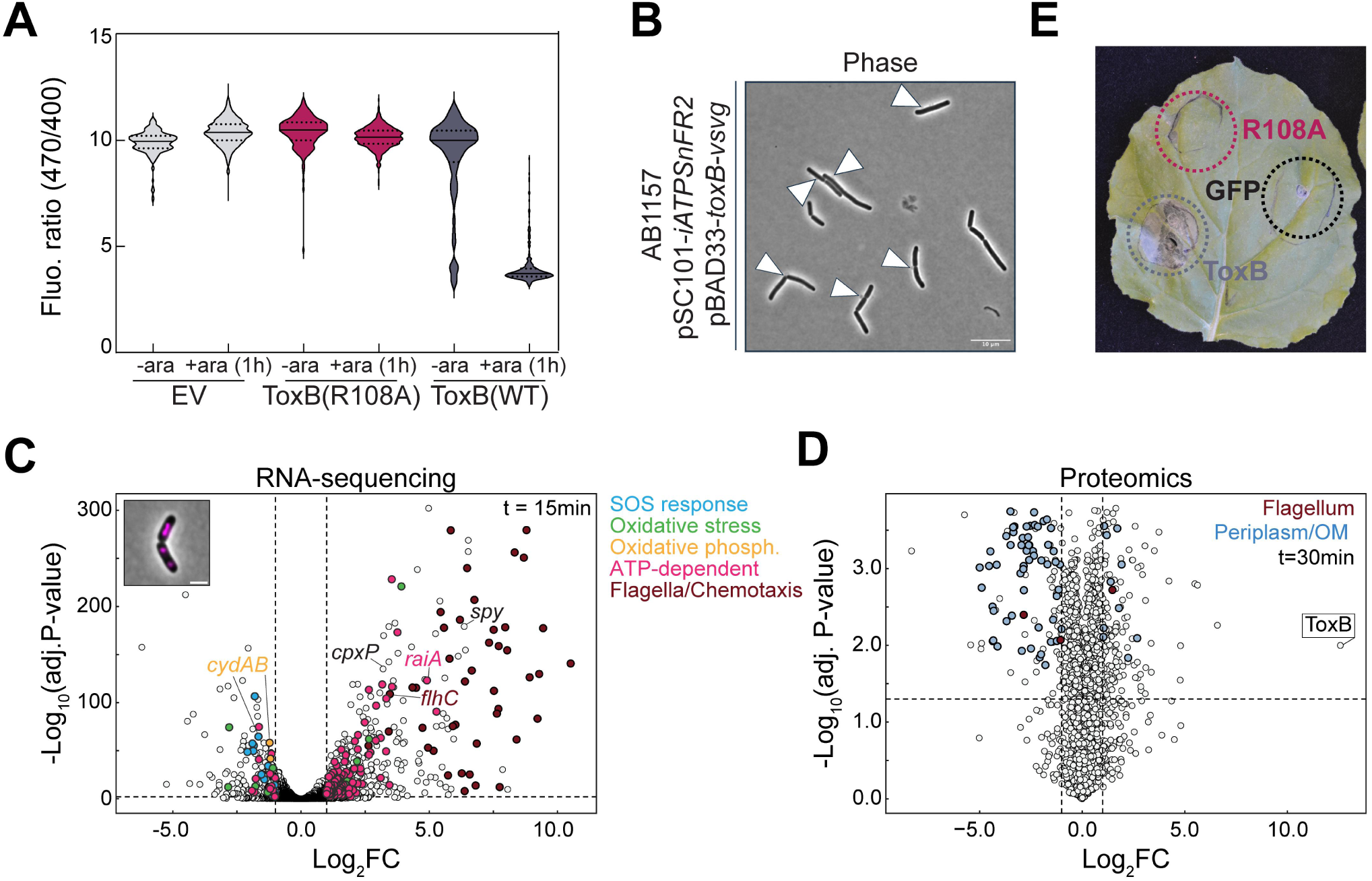
Multi-omics and reporter analyses reveal early metabolic and envelope responses to ToxB intoxication. **(A)** Intracellular ATP measurements using a genetically encoded ATP sensor. Violin plots show single-cell fluorescence ratio values for *E. coli* MG1655 cells carrying empty vector (EV, grey), the catalytic mutant ToxB(R108A) (magenta), or wild-type ToxB (WT, dark purple) under untreated (UT) and induced (1 h) conditions. Each dot represents an individual cell; medians and quartiles are indicated. **(B)** Representative phase-contrast image of intoxicated *E. coli* AB1157 cells corresponding to the experiment in panel A. Arrowheads indicate cells undergoing cell-wall rupture. Scale bar, 10 μm. **(C)** RNA-seq volcano plot showing differential gene expression in *E. coli* AB1157 cells 15 min after induction of *toxB* (WT) compared with empty vector controls. The x-axis shows log₂ fold change (Log₂FC) and the y-axis shows-Log₁₀ adjusted *P* values. Vertical dashed lines indicate Log₂FC thresholds. Genes associated with the SOS response (blue), oxidative stress (green), oxidative phosphorylation (orange), ATP-dependent processes (pink), and flagellar/chemotaxis functions (red) are highlighted. Three independent biological replicates were performed. **(D)** Proteomics volcano plot showing differential protein abundance in ToxB(WT)-expressing cells compared with empty vector controls at 30 min after induction. The x-axis shows Log₂FC and the y-axis shows −Log₁₀ adjusted *P* values. Proteins predicted to localize to the periplasm are highlighted in blue; flagellar proteins are highlighted in red. **(E)** Transient expression assay in *Nicotiana benthamiana* leaves. Agrobacterium-mediated infiltration was used to express GFP, wild-type ToxB, or catalytic mutant ToxB(R108A). Representative leaf images are shown at the indicated time point after infiltration, illustrating differential tissue responses. See also Figures S7-8 and Supplementary Tables 2-5.

To dissect the temporal order of cellular failure and capture early responses prior to cell lysis, we combined RNA-seq at 15 and 30 minutes with quantitative proteomics. Extensive transcriptional remodeling was already evident at 15 minutes (**Figure 8C, Figure S7A-C, Supplementary Tables 2-3**). Gene ontology analysis (**Figure S7A**) revealed perturbation of respiratory metabolism, oxidative phosphorylation (**Figure 8C**, genes highlighted in orange in the volcano plot), and ATP-dependent cellular processes (**Figure 8C**, genes highlighted in pink in the volcano plot), alongside enrichment of stress-associated pathways, including oxidative stress (**Figure 8C**, genes highlighted in green in the volcano plot) and envelope-related responses (**Figure S7A**). Notably, transcripts encoding subunits of the cytochrome *bd* oxidase complex (*cydA* and *cydB*), a terminal enzyme of the bacterial respiratory chain, were significantly downregulated (**Figure 8C, Figure S7C**, genes highlighted in orange in the volcano plot). Beyond its role as a terminal electron acceptor, cytochrome *bd* oxidase also functions as a reactive oxygen species scavenger^67^, and its early suppression suggests that ToxB production compromised oxidative stress defenses. Indeed, oxidative stress genes were found on both sides of the volcano plot (**Figure 8C, Figure S7C**, genes highlighted in green), with primary defense genes, including *katG* and *ahpC*, downregulated and stress-response genes, including *sodA* and *katE*, induced (**Supplementary Tables 2-3**)^68^. Together with the early CellROX signal detected by live-cell imaging (**Figure 6G**), these data indicate that oxidative stress is triggered within the first 15 minutes of ToxB production. Despite the early oxidative stress, the SOS response was not induced but was instead actively repressed: 10 canonical SOS genes, including *recA, dinB, uvrA, uvrB, sulA, umuC, umuD, recN, dinI,* and *yebG*, were all significantly downregulated at 15 minutes and remained so at 30 minutes (**Figure 8C, Figure S7C**, genes highlighted in blue, **Supplementary Tables 2-3**). This is consistent with the lack of induction of the P*_recA_-mScarlet* reporter, and with intact genomic DNA detected by gel electrophoresis (**Figure 6F, Figure S4F**), together supporting the conclusion that ToxB-induced defects do not trigger the canonical SOS response. At the same time, *raiA*, encoding ribosome-associated inhibitor A^69,70^, was among the most strongly upregulated transcripts at this early timepoint (**Figure 8C, Supplementary Table 2**).

By 30 minutes, transcriptional perturbation extended to processes linked to ribosome assembly, translation, and RNA modification, indicating broader metabolic disruptions (**Figure S7B, Supplementary Table 2-3**). Key energetic genes, including the ATP synthase subunit *atpD,* were increasingly repressed, and transcripts for envelope stress markers *cpxP* and *spy* continued to rise (**Figure 8C, Supplementary Tables 2-3**), indicating that the early specific disruptions give way to a broader metabolic and structural collapse. RNA-seq showed transcriptional enrichment of motility-related genes, including essentially all components of the flagellar regulon (**Figure 8C, Figure S7C** genes highlighted in red in the volcano plots, **Figure S7A-B, Supplementary Tables 2-3**); however, quantitative proteomics did not detect increased abundance of flagellar proteins (**Figure 8D, Figure S8A**, genes highlighted in red in the volcano plots, **Supplementary Tables 4-5**). This mRNA-protein disparity was most clearly illustrated by the master flagellar regulator FlhC: its transcript remained significantly upregulated at both timepoints (**Figure 8C, Supplementary Table 2**), yet FlhC protein was among the most strongly depleted at 60 minutes (**Figure S8A, Supplementary Table 4**), consistent with the early translational shutdown evidenced by *raiA* induction. Moreover, proteomic analysis showed marked depletion of numerous periplasmic and outer membrane proteins ∼1h post-induction (**Figure S8A**; genes highlighted in blue in the volcano plot, **Supplementary Table 5**). Given their localization, their apparent loss likely reflects leakage of periplasmic contents following cell envelope disruption rather than transcriptional repression. This interpretation is consistent with microscopy data showing membrane permeabilization at later stages of ToxB production and with periplasmic labelling experiments, in which we fluorescently labelled the periplasm of ToxB-producing cells with a monomeric superfolded Turquoise2 variant adapted for periplasmic fluorescence (msfTq2^C70V^)^71^, thereby confirming loss of compartmentalization (**Figure S8B**). By contrast, envelope stress proteins, including chaperones Spy and CpxP, were significantly enriched at 60 minutes (**Figure S8A, Supplementary Table 4**), consistent with activation of envelope stress responses and increased demand for protein folding and repair within the cell envelope^72–76^.

Together, these findings suggest a model in which respiratory metabolism, oxidative stress defense, and translational control are disrupted within the first 15 minutes, followed by broader metabolic collapse by 30 minutes and eventual cell envelope disintegration by 60 minutes, preceding the decline in intracellular ATP.

Lastly, the breadth of these effects prompted us to ask whether ToxB targets bacteria-specific processes or conserved cellular functions across domains of life. To address this, we transiently expressed *toxB(WT)* and *toxB(R108A)* in tobacco plants, *Nicotiana benthamiana*. Strikingly, expression of *toxB(WT)* induced localized cell death at infiltration sites, characteristic of a hypersensitive response^77^, whereas the ATP-binding-defective R108A mutant or GFP alone had no effect (**Figure 8E**). This cross-kingdom activity suggests that ToxB does not rely on bacteria-specific targets but instead disrupts conserved cellular processes.

## Discussion

### Physiological consequences of ToxB ATPase activity

The cellular effects of ToxB cannot be explained by simple ATP depletion. Intracellular ATP levels remain largely stable during the early stages of ToxB production and decline only later, after substantial cellular damage has already occurred, arguing against global energy collapse as the primary cause of toxicity. Instead, the earliest transcriptional responses point to something more specific: the simultaneous suppression of distinct cellular defenses within the first 15 minutes (**Figure S7A, Supplementary Tables 2-3**). Within the first 15 minutes, genes involved in oxidative stress protection and iron-sulfur cluster repair are downregulated, including the cytochrome *bd* oxidase complex and *ytfE* (**Figure S7A, Supplementary Tables 2-3**). These changes are expected to increase cellular vulnerability by promoting the accumulation of reactive oxygen species and preventing the repair of damaged cofactors. ToxB therefore appears to destabilize cellular homeostasis well before bulk ATP levels decline.

Although we showed that ATP binding is essential for ToxB toxicity, its exact mechanism of action remains unresolved. There is a possibility that ToxB *in vivo* uses ATP to generate a toxic nucleotide-derived molecule rather than simply hydrolyzing ATP. Several anti-phage and antiviral defense systems employ nucleotide-metabolizing enzymes to produce signaling molecules. For example, toxins synthesize nucleotide alarmones such as ppApp, or TIR-domain enzymes that produce ADP-ribose derivatives, including histidine-ADPR, as immune signals in bacteria^57,58^. Here, we tested whether ToxB could generate ATP-derived products similar to those observed. However, *in vitro* biochemical assays with chromatography detected only ATP-to-ADP conversion, with no evidence for the formation of modified nucleotide species (**Figure 5E, Figure S3B**). Nonetheless, we cannot exclude the possibility that additional cofactors or cellular components present *in vivo* enable the formation of such an ATP-derived product, responsible for ToxB’s toxicity.

### ToxB likely disrupts multiple conserved cellular processes

If ToxB does not act by depleting ATP or generating a toxic nucleotide product, its phenotype is more consistent with interference with essential cellular processes. In line with this hypothesis, ToxB causes strong nucleoid compaction without detectable DNA damage or SOS induction, arguing against a classical genotoxic mechanism and instead pointing to disruption of chromosome organization. Because nucleoid architecture depends on multiple ATP-dependent systems, including SMC complexes such as MukBEF, topoisomerases, and nucleoid-associated proteins, impairment of energy-dependent processes could compromise chromosome organization across several pathways^78–85^. Moreover, the failure to isolate spontaneous suppressor mutants further argues against a single dispensable target. Finally, ToxB showed a phytotoxic effect in an ATP-binding-dependent manner. Whatever ToxB targets must therefore be present and functionally relevant in both bacterial and plant cells. Together, these observations support the idea that ToxB perturbs multiple conserved cellular processes rather than acting through a single discrete target.

### An ancient chromosome-maintenance fold repurposed for chromosome disruption

This broad and pleiotropic phenotype is especially striking considering the evolutionary origin of ToxB. ToxB illustrates how a conserved nucleotide-binding fold can be redirected toward toxic activity. Canonical ParB proteins bind CTP and use nucleotide-driven conformational switching to mediate chromosome segregation. In canonical ParB, CTP binding promotes closure of an N-terminal gate to form a DNA-sliding clamp, while the C-dimerization domain provides the constitutive dimerization interface required for this cycle^18^. Recent ancestral reconstruction suggests that such CTP-dependent regulation is an ancient feature of ParB proteins: a reconstructed ParB ancestor from early bacterial evolution already bound *parS* DNA and used CTP-dependent clamp loading, much like modern canonical ParB^15^. ToxB therefore appears to represent a later functional divergence of this ancestral CTP-regulated scaffold toward ATP-dependent toxicity. Its structure retains the conserved phosphate-binding core of the fold, corresponding to Box I (C-motif) and Box II (P-motif), but these motifs are remodeled to favor ATP over CTP. At the same time, ToxB lacks key structural elements required for canonical ParB switching, including the C-terminal dimerization domain and Box III, and therefore is not organized to form the dimeric DNA clamp used by canonical ParB^14^. Together, these features indicate that the ParB-CTPase fold has been repurposed from a CTP-dependent regulatory switch to a monomeric ATP-dependent enzymatic effector.

This type of repurposing is not unique to ToxB. In other nucleotide-binding fold families, evolutionary diversification has repeatedly altered either ligand preference, biochemical output, or biological role. For example, P-loop NTPases include both NTP-dependent switches, such as Ras-like GTPases, and NTP-hydrolyzing enzymes that couple nucleotide turnover to mechanical work, such as helicases and ABC transporters^86–88^. In some cases, shifts in biochemical role are accompanied by changes in nucleotide preference. For example, the bacterial sporulation protein SpoIVA appears to have evolved from a TRAFAC-class P-loop GTPase, yet it hydrolyzes ATP to drive irreversible polymerization^86,88,89^. Similarly, within the sirtuin family, the Rossman NAD-binding fold can support either housekeeping metabolic functions. CobB proteins are NAD-dependent housekeeping enzymes involved in protein deacylation and metabolic regulation, whereas related Sir2-domain proteins in anti-phage systems act as toxic NAD-consuming effectors that trigger abortive infection or growth arrest^58,90–98^. ToxB extends this principle to the ParB-CTPase fold^14^, showing that a scaffold ancestrally linked to chromosome segregation and omnipresent across the tree of life can be redirected toward ATP-dependent toxicity.

## Material and Methods

### Homology survey of ParB-CTPase fold-containing proteins

Seed sequences for ToxB were obtained from the conserved domain annotations CDD cd16392 and InterPro IPR039380. Seed sequences were manually trimmed to match the boundaries of the CDD cd16392 domain, aligned, and used to build a hidden Markov model (HMM) with hmmbuild from HMMER v3.4. The resulting HMM was used to search a database of 68,709 prokaryotic genomes^99^ using hmmsearch with default parameters. For subsequent analyses, we retained only ToxB hits with an HMM-aligned region length greater than 50 amino acids. The taxonomic distribution of ToxB homologs was mapped at the bacterial order level onto a GTDB species tree pruned to the order level^14^.

### Protein alignment, sequence logos, and phylogenetic trees

All ParB and homologous sequences were obtained from our previous work^14^. Multiple sequence alignments were generated with MAFFT v7.490 using the L-INS-i algorithm, except for the tree shown in Fig. 2A, for which the ToxB alignment was added to the previous ParB-C domain alignment using mafft--add. Owing to computational constraints, the domain-level phylogeny was inferred with FastTree v2.1.11 using default settings, including the BLOSUM45 substitution matrix; branch support values correspond to SH-like local supports. The resulting tree was midpoint rooted. The ToxB domain phylogeny was inferred with IQ-TREE v2.3.1 using-alrt 1000-nt AUTO-mem 32G - m TEST and rooted using *Bacillus subtilis* ParB as outgroup. Syntenic context was retrieved using GeneSpy. Sequence logos were generated in R with the ggseqlogo package.

### Strains, media, and growth conditions

All strains used or generated in this study are listed in **Supplementary Table 6**. *E. coli* strains were grown in lysogeny broth (LB) or minimal (M9) medium supplemented with 0.2% casaminoacids. Where appropriate, the media were supplemented with antibiotics at the following concentrations (liquid/solid (μg ml^−1^)): carbenicillin (50/100), chloramphenicol (20/30), and kanamycin (30/50).

### Plasmid construction

Plasmids, gBlocks, and oligos generated or used in this study are listed in **Supplementary Tables 7-8**, respectively. For plasmid construction, a double-stranded DNA fragment containing the desired sequence was chemically synthesized (gBlocks, IDT). The target plasmid was double-digested with restriction enzymes and gel-purified. A 10 μl reaction mixture was assembled with 5 μl of 2x Gibson master mix (NEB) and 5 μl of a combined equimolar mixture of purified backbone and gBlock(s). The mixture was incubated at 50°C for 60 min. Gibson assembly was possible owing to sequence homology shared between the digested backbone and the gBlock fragment(s). All resulting plasmids were verified by either Sanger sequencing (Genewiz) or whole-plasmid sequencing (Plasmidsaurus). Plasmids were introduced into *E. coli* DH5α, AB1157, or *E. coli* BL21 pLysS via heat shock transformation.

### Toxicity assays

*E. coli* AB1157 cells were transformed with appropriate plasmids. For spotting, growth curve, arabinose titration, and immunity-rescuing assays, overnight cultures of appropriate strains were diluted to 10^6^ in tenfold increments, and each dilution was spotted onto LB agar plates containing 0.4% glucose or 0.2% arabinose, supplemented with the appropriate antibiotics. Photographs were taken after overnight growth at 37°C. To compare toxicity among the four ToxB proteins, *E. coli* AB1157 strains harboring corresponding plasmids were grown in LB at 37°C overnight under appropriate antibiotic selection. Stationary-phase overnight cultures were diluted to an OD_600_ of 0.01 in fresh LB medium and grown at 37°C with shaking. At OD_600_ of 0.3, 1 ml culture was retrieved and chilled in ice water, and the remaining culture was treated with 0.2% arabinose or 0.4% glucose. At the indicated timepoints post-induction, 1 ml culture was withdrawn, chilled in ice water for 2 min, then cells were pelleted at 13,000 rpm at 4°C and resuspended in ice-cold fresh LB without inducer and kept on ice. These samples were diluted to 10^6^ in tenfold increments, and 4 μl from each dilution was spotted onto LB agar plates containing appropriate antibiotics.

### Growth inhibition assays

For bacterial competition assays, *E. coli* MG1655 carrying pCH1004-VENN-YciB-ToxB-iToxB (Amp^R^) was used as the attacker strain, and *E. coli* MG1655 *hupA-gfp* (Kan^R^) was used as the target strain. Each strain was grown overnight at 37°C in LB medium supplemented with the appropriate antibiotic. Overnight cultures were diluted into fresh LB medium to an OD_600_ of 0.05 and grown at 37°C with aeration to mid-log phase (OD_600_ < 0.3) in the absence of antibiotics. Attacker and target cells were then mixed at an attacker-to-target ratio of 10:1 and incubated for 2 h at 37°C with aeration, as in^43^. Following co-incubation, cultures were serially diluted in LB and plated on LB agar containing kanamycin only to enumerate surviving target cells. In parallel, after the same 2 h co-incubation, mixed cultures were transferred to 96-well plates containing LB supplemented with kanamycin only, and target outgrowth was monitored by measuring OD_600_ in a plate reader.

### Microfluidics

Stationary-phase overnight cultures were diluted to an OD_600_ of 0.01 in fresh LB medium and grown at 37°C with shaking until an OD_600_ of 0.2; afterward, cells were loaded onto B04A microfluidic perfusion plates (CellASIC). We performed medium manipulation using the ONIX microfluidics platform (CellASIC). Cells were first flushed with LB (without inducer) for 3 min, then with LB supplemented with 0.2% arabinose for 7 hours, before being shifted back to LB. We performed medium exchange at 2 psi for each solution. Images were acquired every 5 minutes.

### Live-cell imaging

Stationary-phase overnight cultures of *E. coli* were diluted to an OD_600_ of 0.01 in fresh LB medium and grown at 37°C with shaking until an OD_600_ of 0.2. At this point, they were treated with 0.2% arabinose or 0.4% glucose and then spotted onto 1.2% agarose pads prepared in LB. For time-lapse experiments, the same fields of view on the pad were imaged at the indicated intervals, with the enclosure temperature set to either 28°C or 30°C. In time-course experiments, samples from the induced cultures were collected at regular intervals as indicated and directly spotted onto agarose pads for snapshots. For nucleoid staining experiments, cells were incubated for 5 min prior to imaging, then stained with DAPI at a final concentration of 5 mg/ml. For monitoring oxidative stress, cells were stained with CellRox Green (Thermo Fisher) at a final concentration of 5 µM, incubated in the dark for 15 min, and washed 3 times in PBS before imaging. For PI staining, cells were incubated at a final concentration of 15 μM PI for 15 min in the dark and then immediately imaged on agarose pads.

### Time-course imaging using the ATP sensor

AB1157 *E. coli* cells containing pSC101-*iATPSnFR2* and pBAD33, pBAD33-*toxB-vsvg,* or pBAD33-*toxB(R108A)-vsvg* plasmid were grown overnight in LB supplemented with 0.4% glucose (and appropriate antibiotics). On the following day, cells were diluted to an OD_600_ of 0.01 in LB supplemented with 0.4% glucose (and appropriate antibiotics) and grown at 37°C to an OD_600_ of 0.2-0.3. Then, the untreated samples were collected, and the remaining cultures were centrifuged at 4,000 rpm for 5 min. The supernatant was replaced with LB containing 0.4% arabinose (with the appropriate antibiotics), and image acquisition was performed by spotting cells onto 1% LB agarose pads after an hour.

### Image acquisition

For all microscopy experiments except those using the ATP sensor, images were acquired using a Zeiss Axio Observer microscope equipped with a 100× oil-immersion objective, an HXP 120 V metal halide lamp, and a Hamamatsu ORCA-Flash4.0 digital camera. ATP sensor experiments were performed on a Zeiss Axio Observer microscope equipped with a 100×/1.45 NA Plan Apo Lambda oil objective, an Orca Flash4 V2 CMOS camera (Hamamatsu), and a Colibri 7 LED light source (Carl Zeiss, Germany). Image acquisition was performed using ZEN Blue 3.2 software (Carl Zeiss, Germany). Fluorescence images were acquired using the following excitation and emission settings. Cells containing a P*_recA_*-*mScarlet* reporter were imaged using a 555 nm LED and 00 Nil Red filter set (530–585 nm excitation, 615LP emission) with 20% LED power and 150 ms exposure. Cells containing the iATPsnFR2 sensor were imaged using dual excitation channels: 475 nm LED with a 38HE filter set (450-490 nm excitation, 500-550 nm emission) at 30% LED power and 100 ms exposure, and 430 nm LED (405-420 nm excitation, 535-550 nm emission) at 20% LED power and 80 ms exposure.

### Image processing and figures preparation

Opening and visualization of original microscopy images were performed using the open-source ImageJ/Fiji software^100^, with contrast and brightness settings kept constant across all regions of interest in each figure unless otherwise specified. Where required, channels were registered to correct for drift using the HyperStackReg plug-in for Fiji. Figures were assembled and annotated using Adobe Illustrator. *E. coli* outlines were obtained using Oufti^101^.

### Cell, nucleoid, and spot detection from microscopy images

The automated cell detection tool Oufti was used to detect outlines of *E*. *coli* cells from phase contrast images. Fluorescence signals were added to cell meshes after background subtraction. Oufti was also used to detect diffraction-limited nucleoids with subpixel precision from fluorescence images, using the objectDetection modules. The detected objects were added to the corresponding cell in the Oufti cell lists, including features related to coordinates, morphology, and intensity. The same optimized nucleoid detection parameters were used to ensure consistency and enable comparisons, as previously described^102,103^.

### Quantitative image analysis from cell meshes

Fluorescence-related analysis, nucleoids information, and other properties of individual cells based on microscopy images were extracted from Oufti cellLists data and plotted using custom codes in MATLAB (MathWorks), as described below.

### Fluorescence profiles

The custom Matlab script MeanIntProfile.m, as in^59,102^ was used to obtain mean relative fluorescence profiles. Briefly, the fluorescence profile of each cell (corresponding to the array of fluorescence intensity per cell segment provided by the relevant signal field in the Oufti cellList) was first normalized by the corresponding array of *steparea* values (corresponding to the area of each segment of the cell), then divided by their sum to obtain relative fluorescence values for each cell (to account for potential concentration differences between cells). When needed, arrays of relative fluorescence were oriented based on the position of the maximal fluorescence intensity of the indicated signal in each cell half. Cell length vectors were normalized from 0 to 1, and the corresponding relative fluorescence profiles from individual cells were interpolated to a fixed-dimension vector and concatenated before averaging.

### Fluorescence intensity analysis

The custom Matlab script MeanFluoPerCell.m as in^59,102^ was used to quantify the distribution of mean fluorescence intensity per cell. Briefly, for each cell in the Oufti cellList, the total fluorescence signal (corresponding to the sum of fluorescence intensity values across all cell segments provided by the relevant signal field) was divided by the total cell area to obtain a mean fluorescence intensity value for that cell. Mean fluorescence values were compiled across all cells and frames to generate population-level distributions. Calculations from MATLAB were used to generate final plots in GraphPad Prism v. 9.5.1.

### Kymographs and demographs

Demographs of relative fluorescence intensity in cells sorted by length were plotted as in^102^. When needed, arrays of relative fluorescence intensity values were oriented based on the position of the maximal fluorescence intensity of the indicated signal in each cell half. Kymographs were obtained using the built-in kymograph function in Oufti.

### Nucleoid size measurement

We used the nucleoid area obtained after running the objectDetection module in Oufti as a proxy for nucleoid size, following^103^, who showed that nucleoid area measurements are unaffected by variations in DAPI signal intensity using a similar detection method. The nucleoid area is provided by the object. area field in the Oufti cell lists, and values were collected in MATLAB for all cells, and converted to µm². Calculations of the nucleoid area and nucleoid/cell (NC) ratios from MATLAB were used to generate final plots in GraphPad Prism v. 9.5.1.

## Statistical analyses

The sample sizes and the number of repeats are included in the figure legends. Means, standard deviations, and significances were calculated in GraphPad Prism v. 9.5.1, MATLAB (MathWorks), or R.

### Genomic DNA extraction

To extract total genomic DNA from intoxicated *E. coli* strains, 3 ml of induced cells was collected at regular intervals, centrifuged at 17,000 × *g* for 1 min, and the pellet was resuspended in 300 μl of Cell Lysis Solution (PureGene, Qiagen). Resuspended cells were incubated at 50 °C for 15 min, mixed with 2 μl of RNase A (NEB 20 mg/ml stock concentration), and incubated at 37 °C for 1 h. Samples were cooled to room temperature, mixed with 100 μl of Protein Precipitation Solution (PureGene, Qiagen), vortexed vigorously for 20 s at high speed, left on ice for 10 min, and then centrifuged at 17,000 x *g* for 10 min. The supernatant was added to 600 μl of isopropanol and mixed well by repeated inversion. Samples were centrifuged again at 17,000 x *g* for 1 min, and the supernatant was discarded. Pellets were resuspended in 600 μl of 70% ethanol and centrifuged at 17,000 x *g* for 1 min. The supernatant was discarded, and a final 1 min centrifugation was performed to remove any remaining liquid. The DNA pellet was resuspended in 50 μl of sterile nuclease-free water and incubated at 37°C for 15-30 min. 50 µl of 500 ng of each sample was loaded onto a 1% agarose gel.

### Immunoblots

Stationary-phase overnight cultures were diluted to an OD_600_ of 0.01 in fresh LB medium and grown at 37°C with shaking. At an OD_600_ of 0.3, the culture was induced with 0.2% arabinose or 0.4% glucose. At the indicated time points post-induction, samples were precipitated with TCA as described previously^59^ after being normalized to the final OD_600_. Denatured samples were resolved on 12% Novex WedgeWell gels (Thermo Fisher Scientific) at 190 V for 45 min. Resolved proteins were transferred to a PVDF membrane using the Trans-Blot Turbo Transfer System (BioRad). For α-VSVG immunoblots, the membrane was incubated with a 1:5000 dilution of a monoclonal α-VSVG-peroxidase HRP-conjugated antibody (Abcam). For α-GFP immunoblots, the membrane was incubated with JL-8 monoclonal antibody (Takara), and for α-mCherry immunoblots, with polyclonal mCherry antibody (CAT# PA5-34974, Thermo Fisher). Goat anti-mouse IgG-peroxidase antibody (Sigma) was used as a secondary antibody for JL-8. Goat anti-rabbit IgG-peroxidase antibody (Sigma) was used as a secondary antibody for α-mCherry. Secondary antibodies were diluted according to the manufacturer’s recommendations. Antibody binding was visualized by chemiluminescence from the reaction of horseradish peroxidase with luminol and imaged using an Image Quant LAS 500 camera (GE Healthcare). Figures were prepared using ImageJ and assembled and annotated in Adobe Illustrator.

### Bioinformatics analysis and structural predictions

The tools InterProScan and HHPred were used to annotate protein domains. AlphaFold3 (AF3^104^) was used to generate predicted protein structures. Models were visualized in PyMOL v.2.5.3 (Schrödinger) and prepared for presentation using UCSF ChimeraX v.1.9.

### Protein expression and purification

Proteins used or purified in this study are listed in **Supplementary Table 9**. N-terminally His-tagged 6xHis-ToxB-iToxB complex (WT/mutants) was expressed from plasmid pET21b in *E. coli* Rosetta (BL21 DE3) pLysS competent cells (Merck). To obtain ToxB-mutants, an overnight culture (10 ml) was used to inoculate 1 L of LB containing selective antibiotics. To obtain ToxB(WT), ToxB(R108A) without the immunity partner, used for EnzCheck and MST assays, and ToxB(CORE) for crystallization, an overnight culture (20 ml) was used to inoculate 1 L of LB containing selective antibiotics. For all, the cells were incubated at 37°C with shaking at 220 rpm until an OD_600_ of ∼0.6. Cultures were cooled for 2 h at 4°C before isopropyl-β-d-1-thiogalactopyranoside (IPTG) was added to a final concentration of 0.5 mM. Cultures were incubated overnight at 16 °C with shaking at 220 rpm before cells were pelleted by centrifugation. Cell pellets were resuspended in buffer A (100 mM Tris-HCl, 300 mM NaCl, 10 mM imidazole, 5% (v/v) glycerol, pH 8.0) with 5 mg lysozyme (Merck) and a cOmplete EDTA-free protease inhibitor tablet (Merck) at room temperature for 30 min with gentle rotation. Cells were lysed on ice by 10 cycles of sonication: 15 s on/15 s off at an amplitude of 20 μm and pelleted at 32,000 g for 35 min at 4°C, and the supernatant was filtered through a 0.22 μm sterile filter (Sartorius).

To obtain ToxB mutants (M76A, Y81A, G105A, R108A, W134A), the clear lysate was loaded onto a gravity column containing 2 mL His-Select Cobalt Affinity Gel (Sigma-Aldrich), pre-equilibrated in buffer A. Proteins were eluted from the column using 500 mM imidazole in the same buffer. Purified fractions were analyzed for purity by sodium dodecyl sulfate-polyacrylamide gel electrophoresis (SDS-PAGE). To obtain ToxB(WT) and ToxB_CORE_ clarified lysate was loaded onto a 1 ml HisTrap HP column (Cytiva) pre-equilibrated with buffer A. Protein was eluted from the column using an increasing gradient of imidazole (10–500 mM) in the same buffer. Desired protein fractions were pooled and loaded onto a preparative-grade HiLoad 16/600 Superdex 75 pg column (GE Healthcare) pre-equilibrated with an elution buffer (5% glycerol (v/v), 10 mM Tris-HCl, 100 mM NaCl, pH 8.0). Desired fractions were identified and analyzed for purity by sodium dodecyl sulfate–polyacrylamide gel electrophoresis (SDS-PAGE) before being pooled. For all, aliquots were flash frozen in liquid nitrogen and stored at-80 °C. For protein samples to be used for protein-nucleotide binding MST experiments, Mg^2+^ was introduced via an overnight dialysis step at 4°C in buffer containing 5% glycerol (v/v), 10 mM Tris-HCl, 100 mM NaCl, and 5 mM MgCl_2_, pH 8.0, before concentration and quantification as described above.

### Protein crystallization and X-ray data collection

ToxB-iToxB complex at a concentration of 20 mg/ml in buffer containing 5 % (v/v) glycerol, 10 mM Tris-HCl, and 100 mM NaCl, pH 8.0, was prepared for crystallization by adding 1 mM MgCl2 and 1 mM ATPγS. Commercially available (Molecular Dimensions) crystallization screens in the sitting drop vapour diffusion format were set up in MRC2 96-well crystallization plates (SwissSci) with drops comprised of 0.3 μl precipitant solution and 0.3 μl protein complex using an Oryx 8 liquid handling robot (Douglas Instruments) before being equilibrated against 50 µl reservoir at a constant temperature of 293 K. ToxB-iToxB-ATPγS crystals grew in a solution containing 0.12 M Ethylene glycols; 0.1 M Tris (base); BICINE pH 8.5, 10 % (v/v) PEG 8000 and 20 % (v/v) ethylene glycol. Suitable crystals were mounted in Litholoops (Molecular Dimensions) and flash-cooled by plunging into liquid nitrogen. X-ray data were collected on beamline I04 at the Diamond Light Source (Oxfordshire, UK) using an Eiger2 XE 16M hybrid photon counting detector (Dectris) with crystals maintained at 100 K by a Cryojet cryocooler (Oxford Instruments).

### Structure determination and refinement

Data for the ToxB-iToxB complex were collected from a single crystal at a wavelength of 0.9537 Å and processed using DIALS (version 3.23.0^105^) via the XIA2 expert system. All downstream processing and refinement were performed using the CCP4 Cloud interface (v 1.8.012^106^). Scaled data were merged using AIMLESS (data statistics shown in **Supplementary Table 10**^107^) with a high-resolution cutoff at 2.4 Å. The structure was solved by molecular replacement using PHASER^108^ in space group *P2_1_2_1_2_1_* with cell parameters of *a* = 48.37 Å, *b* = 99.62 Å, *c* = 108.07 Å, and *α* = *β* = *γ* = 90 ° with template models of *Erwinia’s* ToxB and iToxB generated by Alphafold3 (AF3^104^). The resultant model was refined using iterative stages of model building in COOT^109^ and restrained refinement in REFMAC5^110^. The final 3D atomic model was validated using MolProbity^111^ and the PDB-Validation server^112^ prior to deposition.

### Microscale thermophoresis (MST)

MST experiments were performed as follows. ToxB (WT and R108A) was fluorescently labeled using a His-Tag Labeling Kit (RED-tris-NTA 2nd Generation) according to the manufacturer’s protocol. A titration series of ligands (ATP, CTP, UTP, GTP) was generated using a 16-step doubling dilution from 40 mM, and the resulting solutions were mixed in a 1:1 ratio with the fluorescently labeled protein, keeping the final concentration constant at 25 nM. The mixture was incubated at room temperature for 15 minutes before the samples were loaded into Monolith premium capillaries (NanoTemper). MST traces were collected at 100% excitation and medium power using a Monolith X instrument (NanoTemper). The experiment was repeated in triplicate, and the results were plotted and analyzed using GraphPad Prism v. 9.5.1. The thermophoretic mobility of the protein was determined using the normalized fluorescence (Fnorm [‰]) before and after heating. The dissociation constant (K_D_) was calculated using nonlinear regression fitting in GraphPad Prism v. 9.5.1.

### Measurement of ATPase activity by EnzChek phosphate release assay

ATP hydrolysis was monitored using an EnzCheck Phosphate Assay Kit (Thermo Fisher Scientific). Samples (100 μl) containing a reaction buffer supplemented with an increasing concentration of ATP (1, 5, 10, 50, 100, 500, and 1,000 μM), and 1 μM concentration of ToxB (WT or mutants) were assayed in a CLARIOstar Plus plate reader (BMG Labtech) at 25°C for 8 h with readings every 1 min with continuous orbital shaking at 300 rpm between reads. The reaction buffer (1 ml) typically contained 740 μl ultrapure water, 50 μl 20× reaction buffer (100 mM Tris-HCl, 2 M NaCl, and 20 mM MgCl_2_, pH 8.0), 200 μl 2-amino-6-mercapto-7-methylpurine riboside (MESG) substrate solution, and 10 μl purine nucleoside phosphorylase enzyme (one unit). Reactions with buffer only or buffer + NTP were also included as controls. The inorganic phosphate standard curve was also constructed according to the instruction guidelines. The results were analyzed using Excel v. 16.92 and plotted in GraphPad Prism v. 9.5.1. The ATPase rates were calculated using a linear regression fitting in GraphPad Prism v. 9.5.1. Error bars represent standard deviations from three replicates.

### Detection of negatively charged adenosine nucleotide species by SAX-UV

The formation of negatively charged adenosine nucleotide species was followed using strong anion-exchange chromatography coupled with UV detection (SAX-UV). Reactions were performed in a buffer containing 5 mM Tris-HCl, 100 mM NaCl, and 4 mM MgCl_2_ (pH 8.0), with ATP as a substrate. Reactions were initiated by the addition of purified *E. amylovora* ToxB and incubated at room temperature. At the indicated time points, aliquots were collected and quenched by adding an equal volume of methanol. Samples were centrifuged to remove precipitated protein, and the resulting supernatants were analyzed by SAX-UV. Control reactions containing ATP in the buffer without the enzyme were analyzed in parallel. The analysis was performed on a Dionex UltiMate 3000 instrument (Thermo Fisher Scientific) using a Poros HQ 50 strong anion exchange column (10 x 50 mm). Mobile phases used for elution were: A, 5 mM NH_4_HCO_3_; B, 1 M NH_4_HCO_3_. Samples of reaction mixtures or authentic standards (ATP, ADP, AMP) dissolved in water (60 µM) were injected (20 µl), and the column was eluted with a linear gradient of buffer B from 0% to 100% in 10 minutes, held for 4.5 minutes, and then down to 0% B over 1.5 minutes. Finally, the column was equilibrated at 0% B for 4 minutes; all steps were performed at a flow rate of 7 ml per min, and detection was performed with an on-line detector to monitor A_265_. Where necessary, peak identities were confirmed by co-injection with authentic standards. Chromeleon (Dionex, Thermo Fisher Scientific) software was used to process data.

### LC-MS/MS nucleotide identification from purified protein complexes

Ligands bound to purified ToxB-iToxB protein complexes were identified by LC-MS/MS following organic solvent denaturation. Purified protein complexes were mixed 1:1 (v/v) with methanol to denature the proteins and to release any bound small-molecule ligands. Samples were vortexed, centrifuged to pellet precipitated protein, and the supernatant was transferred into a glass insert for analysis. LC-MS/MS analysis was performed on a triple quadrupole mass spectrometer (Xevo TQ Absolute, Waters) coupled to liquid chromatography using a porous graphitic carbon column (Hypercarb, Thermo Scientific) as described previously^113^. In brief, nucleotides were separated using a gradient of ammonium formate buffer adjusted to pH 9.0 and acetonitrile and detected in negative electrospray ionization mode using multiple reaction monitoring (MRM). Standard nucleotide mixtures (ATP, ADP, CTP, CDP, GTP, GDP, UTP, UDP) were analyzed under identical conditions to determine retention times and confirm compound identity. Calibration curves were obtained for ATP, ADP, and CTP using serial dilutions of standards spanning concentrations from 1 to 100 μM. Data were processed using MassLynx software.

### RNA extraction and sequencing

Stationary-phase overnight cultures were diluted to an OD_600_ of 0.01 in fresh LB medium and grown at 37°C with shaking. At OD_600_ of 0.2, cells were incubated with 0.4% glucose or 0.2% arabinose until the indicated time points. 5 ml of cell cultures at each time point were pelleted for RNA extraction. Total RNA was purified using the Direct-zol RNA miniprep kit (Zymo Research), and 10 μg was incubated with 20 units of Turbo DNaseI (Invitrogen) at 37 °C for 1 h to remove any contaminated genomic DNA. DNaseI was subsequently removed using the RNA Clean and Concentrator-25 kit (Zymo Research). Purified RNA samples were processed by Genewiz where bacterial rRNA was depleted using an NEBNext rRNA Depletion Kit (Cat# E7850X). DNA libraries were prepared by Genewiz and sequenced on an Illumina NovaSeq platform. RNA-seq data, consisting of short, paired-end Illumina reads, were analyzed as described below.

### Total cell proteome of cells expressing ToxB

*E. coli AB1157* strain carrying the empty vector pBAD33 and pBAD33-*toxB(Ea)-vsvg* were grown overnight in 3 mL of LB supplemented with 0.4% glucose and chloramphenicol (34 µg/ml), at 37 °C, 220 rpm. The overnight cultures were diluted 30-fold in fresh LB supplemented with chloramphenicol and grown until the OD_600_ reached 0.4. Samples were collected at time 0 h, and the remaining culture was divided into aliquots taken after 30 minutes and 1 hour, both with and without induction using 0.4% arabinose. The aliquots were pelleted and washed twice with cold PBS 1 x, and the pellets were stored at-20 °C until further processing at the Max Planck Institute for Terrestrial Microbiology Mass Spectrometry and Proteomics Facility.

For protein extraction, cell pellets were resuspended in 2% sodium lauroyl sarcosinate (SLS) and 100-mM ammonium bicarbonate. Cells were lysed by incubation at 90 °C for 15 min, followed by sonication (Vial Tweeter, Hielscher) at 80% amplitude for 30 seconds. Cell lysates were reduced by adding 5 mM (final concentration) Tris(2-carboxyethyl)phosphine and incubated at 95 °C for 15 min, followed by alkylation (10-mM iodoacetamide final concentration, 30 min at 25 °C). The amount of extracted proteins was measured using the BCA protein assay (Thermo Fisher Scientific). Fifty microgram total protein was then digested with 1 µg trypsin (Promega) overnight at 30 °C in the presence of 0.5% SLS. Following digestion, SLS was precipitated with trifluoroacetic acid (TFA, 1.5% final concentration), and peptides were purified using Chromabond C18 microspin columns (Macherey-Nagel). Acidified peptides were loaded onto spin columns equilibrated with acetonitrile and then 0.15% TFA. After peptide loading, a washing step with 0.15% TFA was performed, followed by elution using 50% acetonitrile. Eluted peptides were then dried by a vacuum concentrator and reconstituted in 0.15% TFA. Peptide mixtures were analyzed by liquid chromatography-mass spectrometry using an UltiMate 3000 RSLCnano connected to a Q-Exactive Plus mass spectrometer (both Thermo Scien,tific) as reported previously^114–116^. The data were further analyzed using either Progenesis (Waters) or MaxQuant in standard settings^117^ using an *E. coli* Uniprot database. Follow-up data analysis and data visualization were done with SafeQuant^117^ (available under https://github.com/eahrne/SafeQuant), Perseus^118^, and RStudio software. All the analysis steps were done in three biological replicates, and proteins enriched or depleted more than 2-fold and with a p-value lower than 0.05 were highlighted.

### RNA-seq and proteomics data analysis

RNA-seq data were analyzed using the Galaxy Australia platform (https://usegalaxy.org.au/)^119^. Read quality was assessed using FastQC^120^ and summarized with MultiQC^121^. Adapter removal and quality trimming were performed using Trimmomatic^122^, including an initial ILLUMINACLIP step (3:30:10:8), followed by SLIDINGWINDOW trimming (3:10) and removal of reads shorter than 20 bp (MINLEN:20). Trimmed reads were aligned to the *E. coli* reference genome (NCBI RefSeq accession GCF_000005845.2, assembly ASM584v2) using HISAT2^123^. Gene-level read counts were generated with HTSeq^124^. Differential gene expression analysis was performed using DESeq^125^. Genes were considered differentially expressed if the adjusted p-value was < 0.01 and |log₂ fold change| > 1. Downstream data processing and annotation were performed using custom scripts available at https://doi.org/10.5281/zenodo.18781500. Gene Ontology (GO) enrichment analysis was conducted using the GOseq package^126^ in R v4.5.0. Enriched GO categories were visualized as bubble pots showing the top 20 terms using ggplot2^127^ using custom R scripts (https://doi.org/10.5281/zenodo.19705258). Genes without GO annotations were excluded from enrichment analysis. Additional visualizations for RNA-seq and Proteomics data, including volcano plots, were generated using custom Python scripts and notebooks (https://doi.org/10.5281/zenodo.19705556). Genes belonging to individual GO categories were extracted using a custom Python script available at https://doi.org/10.5281/zenodo.18786120. Final plotting and data visualization were performed using Python 3.10 and Matplotlib.

### Plant cell death assay

The cell death assay was performed as previously described^77^. For transient expression experiments, *Agrobacterium tumefaciens* strain GV3101^128^ carrying the respective constructs (**Supplementary Table 6**) was used. To enable cloning and propagation of the toxin gene in *E. coli* while preventing toxicity, intron 1 from the *Arabidopsis thaliana ssp5* gene (AT5G11860) was inserted into the *toxB*(WT) coding sequence and *toxB(R108A)* mutant. The presence of the intron introduces a frameshift in *E. coli*, preventing toxin expression, whereas the intron is correctly spliced in plant cells, restoring the functional coding sequence. *Agrobacterium* strains harbouring plasmids for transient expression of *toxB(WT), toxB(R108A), or gfp* (negative control) were grown for 2 days in 10 mL LB medium supplemented with the appropriate antibiotics. Cells were harvested by centrifugation, washed, and resuspended in infiltration buffer (10 mM MgCl₂, 10 mM MES, 0.2 mM acetosyringone). The suspensions were incubated at room temperature for at least 3 hours before infiltration. *Agrobacterium* suspensions were adjusted to an OD_600_ of 0.3 and spot-infiltrated into leaves of 4-5-week-old *Nicotiana benthamiana* plants. One day after infiltration, protein expression was induced by spraying the leaves with 10 µM dexamethasone and 0.05% Silwet L-77. Cell death phenotypes were assessed 4 days post-infiltration. Three leaves per plant were infiltrated, and three plants were used per assay.

### Data, resources, and software availability

The R codes and datasets have been deposited at Zenodo.

## Author contribution

Conceptualization: J.K, T.B.K.L

Methodology: J.K, J.E.A.M, B.S, M.R, T.O, F.G, T.C.M, V.S, S.E, L.V.M, A.H

Investigation: J.K, J.E.A.M, B.S, M.R, T.O, F.G, T.C.M, K.E.J, A.S.M, N.T.T, S.E, L.V.M, A.H Visualization: J.K, J.E.A.M, B.S, T.O, S.E, A.H

Funding acquisition: V.S, L.V.M, A.H, T.B.K.L

Project administration: J.K, V.S, S.E, L.V.M, A.H, T.B.K.L Supervision: V.S, S.E, L.V.M, A.H, T.B.K.L

Writing – original draft: J.K

Writing – review & editing: J.K, J.E.A.M, B.S, M.R, T.O, F. G, T.C.M, V.S, S.E, L.V.M, A.H, T.B.K.L

## Acknowledgements

We thank members of the Le lab, Géraldine Laloux, Susan Schlimpert, Martin Thanbichler, Mark Buttner, Yoann Santin, and Miloš Tišma for insightful discussions and critical reading of the manuscript. We are grateful to Anjana Badrinarayanan for providing strains, Robert Sablowski for plant infiltration assay resources, and Chris Hayes for strains and valuable advice. We also thank Susan Schlimpert for support with microfluidics experiments, Timo Glatter and Jörg Khant for the support with proteomics analysis. This work is supported by a Lister Institute Fellowship and Wellcome Trust Investigator grant 221776/Z/2/Z (to T.B.K.L.) that funds J.K, T.C.M, and K.E.J., and the BBSRC-funded Harnessing Biosynthesis for Sustainable Food and Health (HBio) Institute Strategic Programme BB/X01097X/1 (to the John Innes Centre that funds N.T.T). We thank Diamond Light Source for access to beamline I04 under proposal MX32728. A.H. is supported by a Wellcome Trust Career Development Award (227755/Z/23/Z).

---

**Movie S1.** ToxB production in *E. coli* cells leads to cell chaining and cell lysis; related to Figure 6A

**Movie S2.** ToxB production in *E. coli* cells with constitutive mCherry cytoplasmic labeling; related to Figure S4A

**Movie S3.** ToxB production in *E. coli* cells expressing HU-GFP; related to Figure S4D

**Movie S4.** ToxB N106A mutant remains toxic in *E. coli* cells; related to Figure S6D

**Supplementary Table 1.** Details of Box I (C motif) and Box II (P motif) nucleotide contact residues in ToxB-ATP protein based on the ToxB-iToxB crystal structure

**Supplementary Table 2.** RNA-seq differential expression analysis of *E. coli* cells expressing ToxB versus empty vector control, including log2 fold change and adjusted p-values

**Supplementary Table 3.** Gene Ontology (GO) enrichment analysis of differentially expressed genes from Supplementary Table 2

**Supplementary Table 4.** Proteomics differential abundance analysis of *E. coli* cells expressing ToxB versus empty vector control, including protein identifiers, fold changes, and adjusted p-values

**Supplementary Table 5.** Gene Ontology (GO) enrichment analysis of differentially abundant proteins (ToxB vs empty vector) from Supplementary Table 4, including log2 fold change and adjusted p-values

**Supplementary Table 6.** Description of bacterial strains used in this study **Supplementary Table 7.** Description of plasmid constructions used in this study **Supplementary Table 8.** Description of oligonucleotides used in this study **Supplementary Table 9.** Description of proteins purified and analyzed in this study

**Supplementary Table 10.** Data collection and refinement statistics for the ToxB–iToxB crystal structure

## Supplementary figure legends

**Figure S1.**
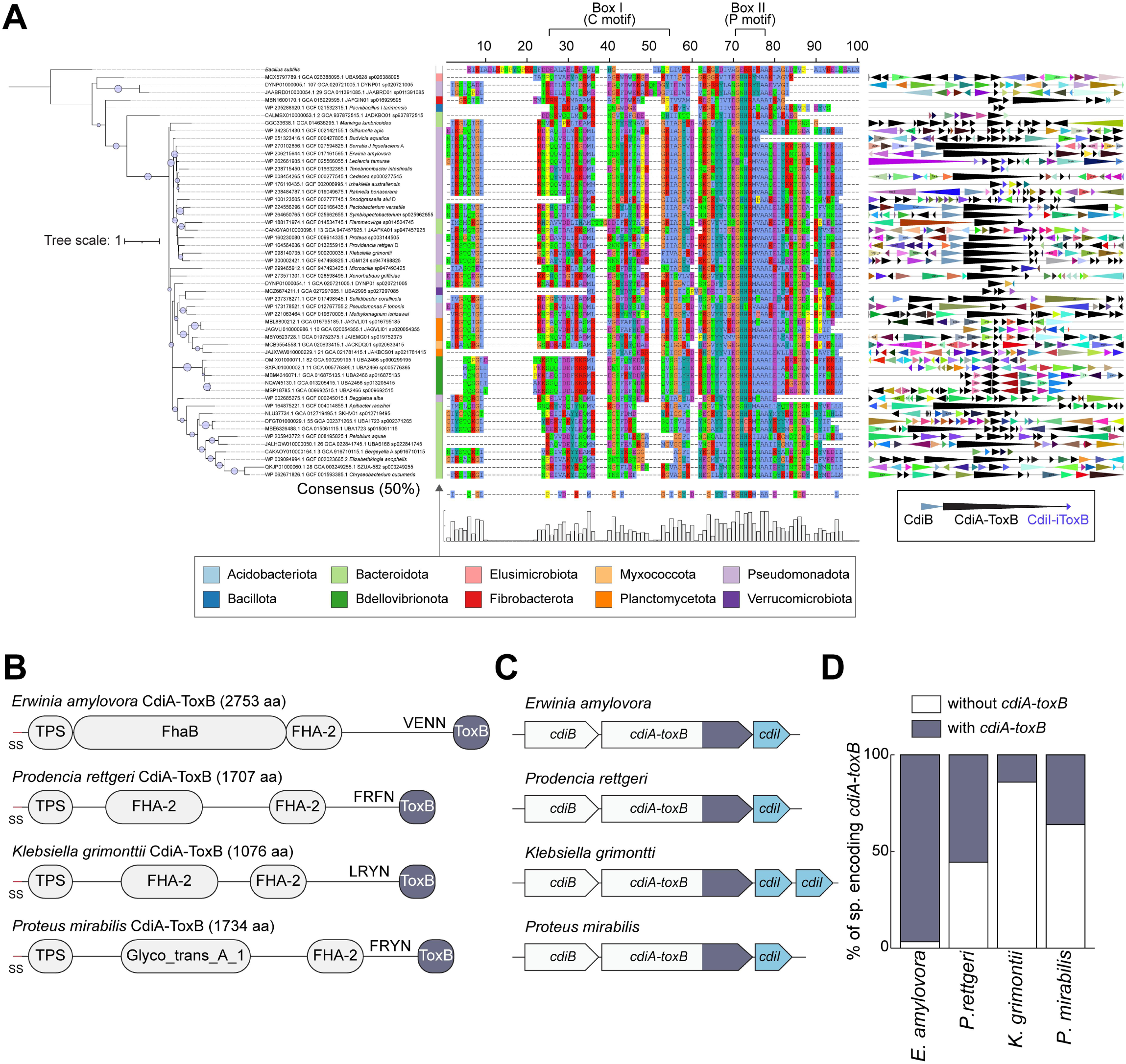
Phylogenetic distribution, sequence conservation, and genomic context of ToxB-containing CDI systems. Related to. **Figure 2. (A)** Phylogenetic tree of ToxB homologs identified across bacterial genomes, colored by phylum. The tree is shown alongside a multiple sequence alignment of the CdiA-ToxB C-terminal domain, with conserved regions corresponding to the ParB-CTPase fold highlighted. Box I (C motif) and Box II (P motif) are indicated above the alignment. Gene neighbourhood schematics on the right show representative *cdiB-cdiA-toxB-cdiI* organizations, including variations in the *toxB* size, number, and arrangement of immunity genes. **(B)** Domain architecture of representative CdiA-ToxB proteins from *Erwinia amylovora*, *Providencia rettgeri*, *Klebsiella grimontii*, and *Proteus mirabilis*. Annotated domains include the N-terminal signal sequence (SS), type Vb secretion system (TPS) domain, FHA repeat regions or other scaffold domains, variable linker motifs, and the conserved C-terminal ToxB toxin domain. **(C)** Genomic organization of CDI loci encoding ToxB in the same representative species. Arrows indicate *cdiB*, *cdiA-toxB*, and downstream *cdiI* immunity genes. Loci with multiple immunity genes are shown. **(D)** Comparison of the total number of CdiA proteins and the subset encoding a ToxB C-terminal domain in the indicated species.

**Figure S2.**
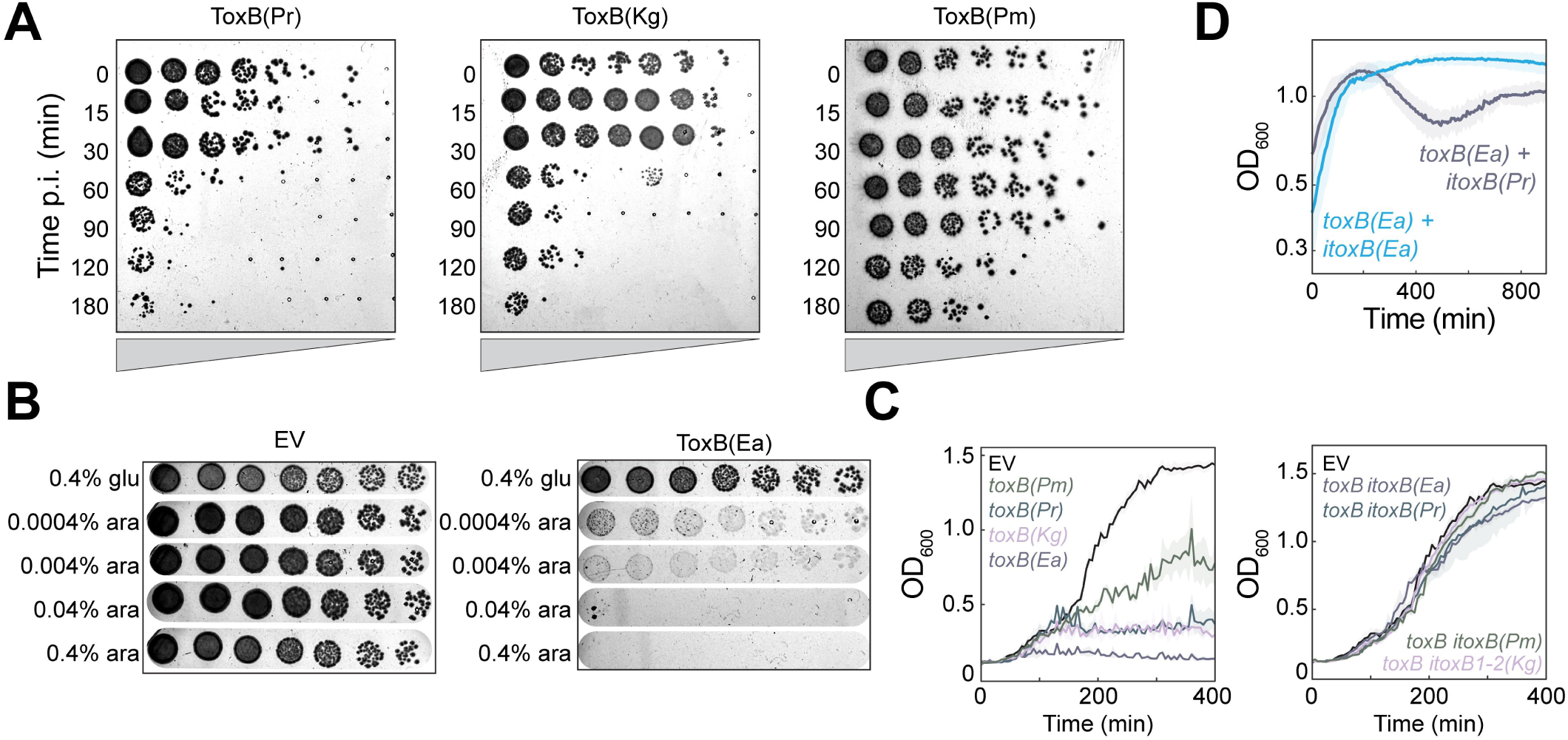
ToxB acts as a bacterial toxin, and iToxB as its immunity partner. Related to. **Figure 3. (A)** Time-course viability assays following induction of ToxB homologs from *Providencia rettgeri* ToxB(Pr), *Klebsiella grimontii* ToxB(Kg), and *Proteus mirabilis* ToxB(Pm). Experiments were performed as in Fig. 3D. **(B)** Spot dilution assays of AB1157 *E. coli* cells carrying an empty vector or pBAD33-*toxB(Ea)* at increasing glucose (left) or arabinose (right) concentrations. **(C)** Growth curves of *E. coli* cells expressing all *toxB* homologs alone (left) or together with their corresponding immunity proteins (right), performed under inducing conditions, as in Fig. 3E. **(D)** Growth curves of *E. coli* cells expressing *toxB(Ea)* together with their corresponding immunity protein, iToxB(Ea) (purple), or with their non-cognate immunity protein, iToxB(Pr) (blue), performed under inducing conditions, as in Fig. 3C.

**Figure S3.**
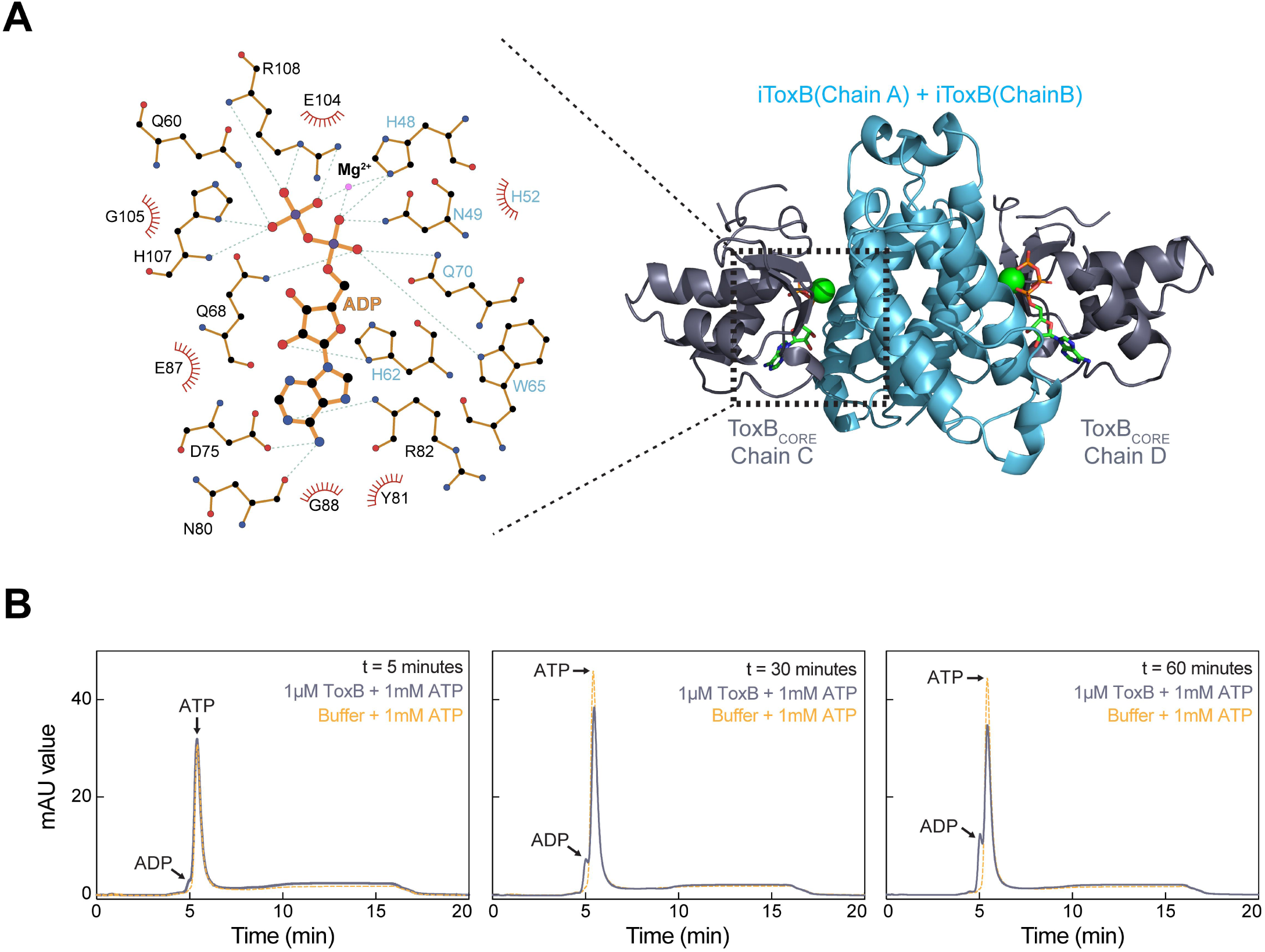
Biochemical characterization of ATP binding and hydrolysis by ToxB. Related to. **Figures 4-5. (A)** LigPlot representation of ADP binding in a symmetry-related ToxB active site. Schematic interaction map showing contacts between ADP and residues from the toxin and the immunity protein. Hydrogen bonds are indicated by green dashed lines and hydrophobic contacts by red arcs. Residues shown in blue originate from the immunity protein (iToxB). Note that the lack of the 𝜸-phosphate leads to an ordering of the N-terminal residues of the ADP-binding ToxB monomer, which are disordered in the ATP-binding ToxB counterpart, and results in residue Q60 adopting the position of the missing 𝜸-phosphate, thereby displacing part of the C-terminus. **(B)** ATP is converted to ADP *in vitro*. As in Figure 5E, only for t = 5, 30, and 60 min. Buffer-only controls are shown for comparison.

**Figure S4.**
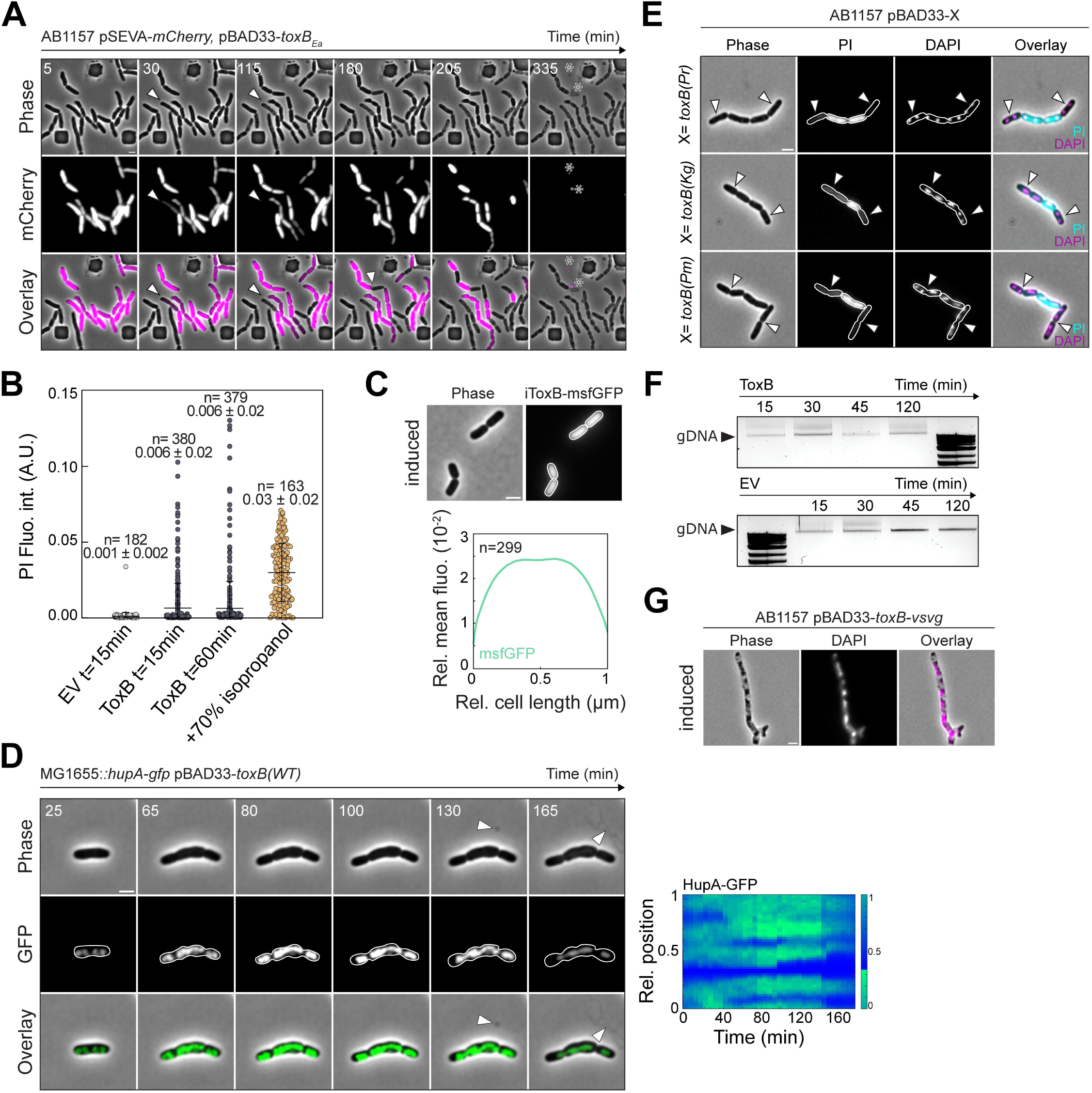
ToxB intoxication leads to late membrane permeabilization without detectable DNA cleavage. Related to. **Figure 6. (A)** Cytoplasmic contents are lost during late-stage intoxication. Time-lapse microfluidics imaging of AB1157 cells expressing *toxB* and constitutively expressing cytoplasmic mCherry from pBAD33-*toxB-vsvg*, and pSEVA251-*mCherry*, respectively. Phase-contrast, mCherry, and overlay images are shown at the indicated times. Arrowheads point towards cells that lost the cytoplasmic mCherry signal; asterisks point towards the mCherry signal outside of the cell. **(B)** Membrane permeabilization increases over time following ToxB induction. Single-cell PI fluorescence intensity measured under repressing and inducing conditions at the indicated times following toxB induction. Cells treated with 70% isopropanol serve as a positive control. Mean ± SD and n values are indicated. **(C)** iToxB localizes to the cytosol. Representative phase-contrast and fluorescence images of AB1157 cells expressing *itoxB-msfgfp* from pBAD33. Right, mean pole-to-pole fluorescence intensity profiles of msfGFP signal. **(D)** ToxB intoxication disrupts chromosome organization prior to envelope failure. Time-lapse imaging of MG1655 *E. coli hupA-gfp frt-kan* cells expressing *toxB* from pBAD33. Phase-contrast, HupA-GFP, and overlay images are shown at the indicated times. Arrowheads indicate compromised cellular envelope and release of cellular content. Right, kymograph of HupA-GFP fluorescence intensity along the cell length for a representative cell. **(E)** ToxB homologs induce membrane permeabilization. Propidium iodide (PI) staining of AB1157 *E. coli* cells expressing ToxB homologs ToxB(Pr), ToxB(Kg), and ToxB(Pm). Representative phase-contrast, DAPI, PI, and overlay images are shown. Arrowheads indicate cells with condensed DNA that are PI-negative, indicating that membrane integrity is still intact. **(F)** ToxB intoxication does not cause detectable genomic DNA cleavage. Genomic DNA isolated from *toxB*-expressing or empty vector control cells at the indicated times after induction was analyzed by agarose gel electrophoresis. **(G)** Chromosome compaction persists at late time points. Representative phase-contrast, DAPI fluorescence, and overlay images of AB1157 cells expressing *toxB* from pBAD33-*toxB-vsvg* 5 h after ToxB induction. Unless otherwise indicated, experiments were performed using *Erwinia amylovora* ToxB(Ea). For all, induction and repression were performed with 0.4% arabinose and 0.4% glucose, respectively. For all, cell outlines were generated using Oufti; scale bars, 2 µm.

**Figure S5.**
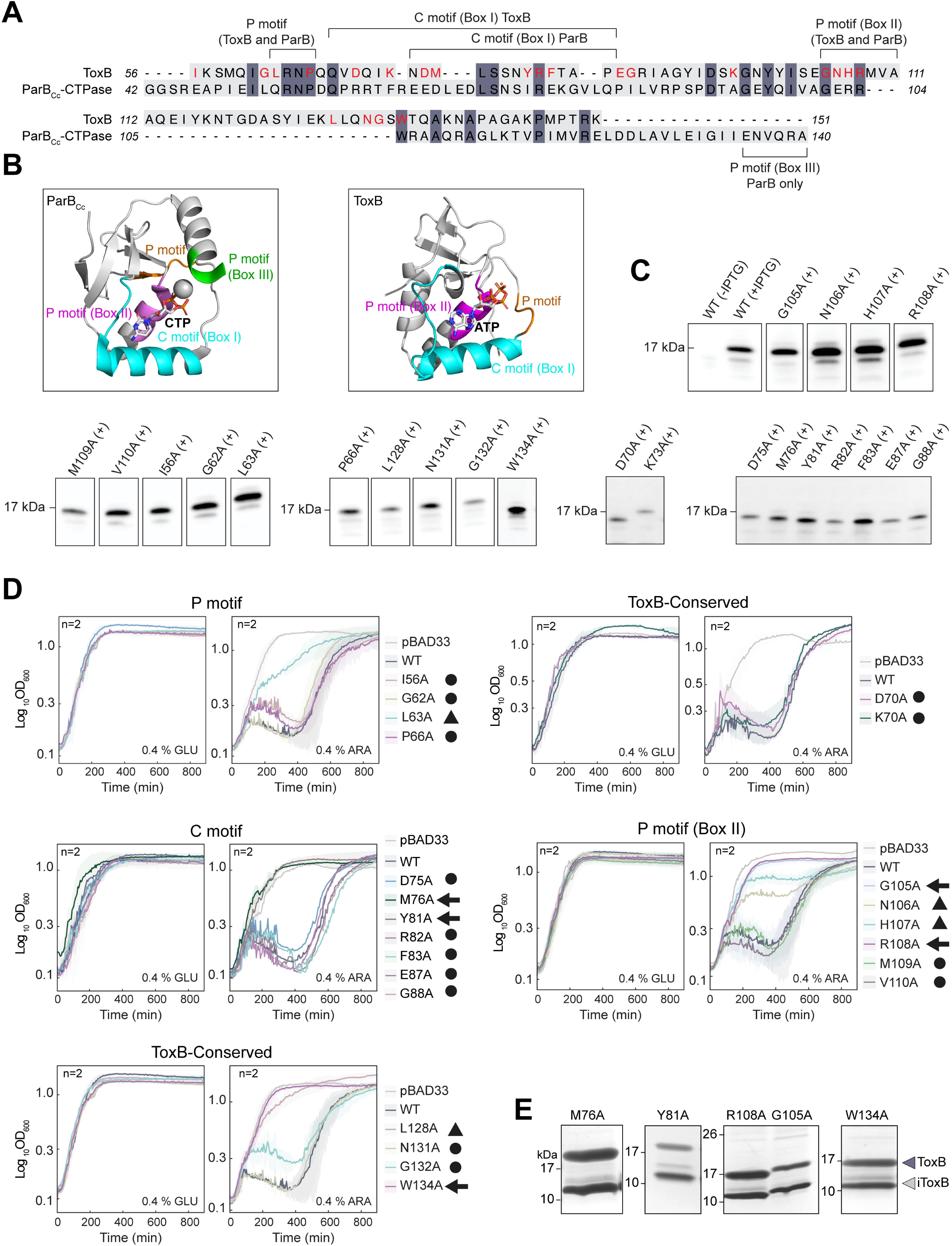
Sequence conservation and functional analysis of the ParB-CTPase motifs in ToxB. Related to. **Figure 7. (A)** Sequence alignment of the ParB-CTPase domain and ToxB, colored by residue identity. Conserved motifs are indicated, including the C motif (Box I) and P motifs (Box II and Box III). Residues selected for alanine mutagenesis in ToxB (red) were chosen based on a combination of sequence conservation across ToxB homologs and structural correspondence to the ParB fold, with particular focus on residues positioned within the nucleotide-binding region. **(B)** Structural comparison of the ParB-CTPase domain and ToxB. Conserved motifs are mapped onto the structures, including the C motif (Box I) and P motif (Box II), and Box III in ParB. Bound nucleotides (CTP in ParB and ATP in ToxB) are shown. Structural differences highlight repositioning of motif elements within the nucleotide-binding pocket. **(C)** ToxB alanine substitution mutants are expressed at comparable levels. Western blot analysis of ToxB alanine substitution mutants. Whole-cell extracts from *E. coli* expressing VSVG-tagged ToxB mutants were probed with an anti-VSVG antibody. **(D)** Specific residues within the conserved ParB-CTPase fold are required for ToxB toxicity. Growth curves of *E. coli* cells expressing ToxB alanine substitution mutants under repressing (0.4% glucose, left) or inducing (0.4% arabinose, right) conditions. Mutants are grouped according to their position within the P and C motifs, or ToxB-conserved regions. Symbols indicate functional class: circles represent wild-type-like toxicity (Class I); triangles, attenuated toxicity (Class II), and arrows, non-toxic mutants (Class III). Experiments were performed in duplicate. **(E)** Class III mutants retain interaction with the immunity protein. Co-immunoprecipitation analysis of representative non-toxic ToxB mutants. Unless otherwise indicated, experiments were performed using *Erwinia amylovora* ToxB(Ea).

**Figure S6.**
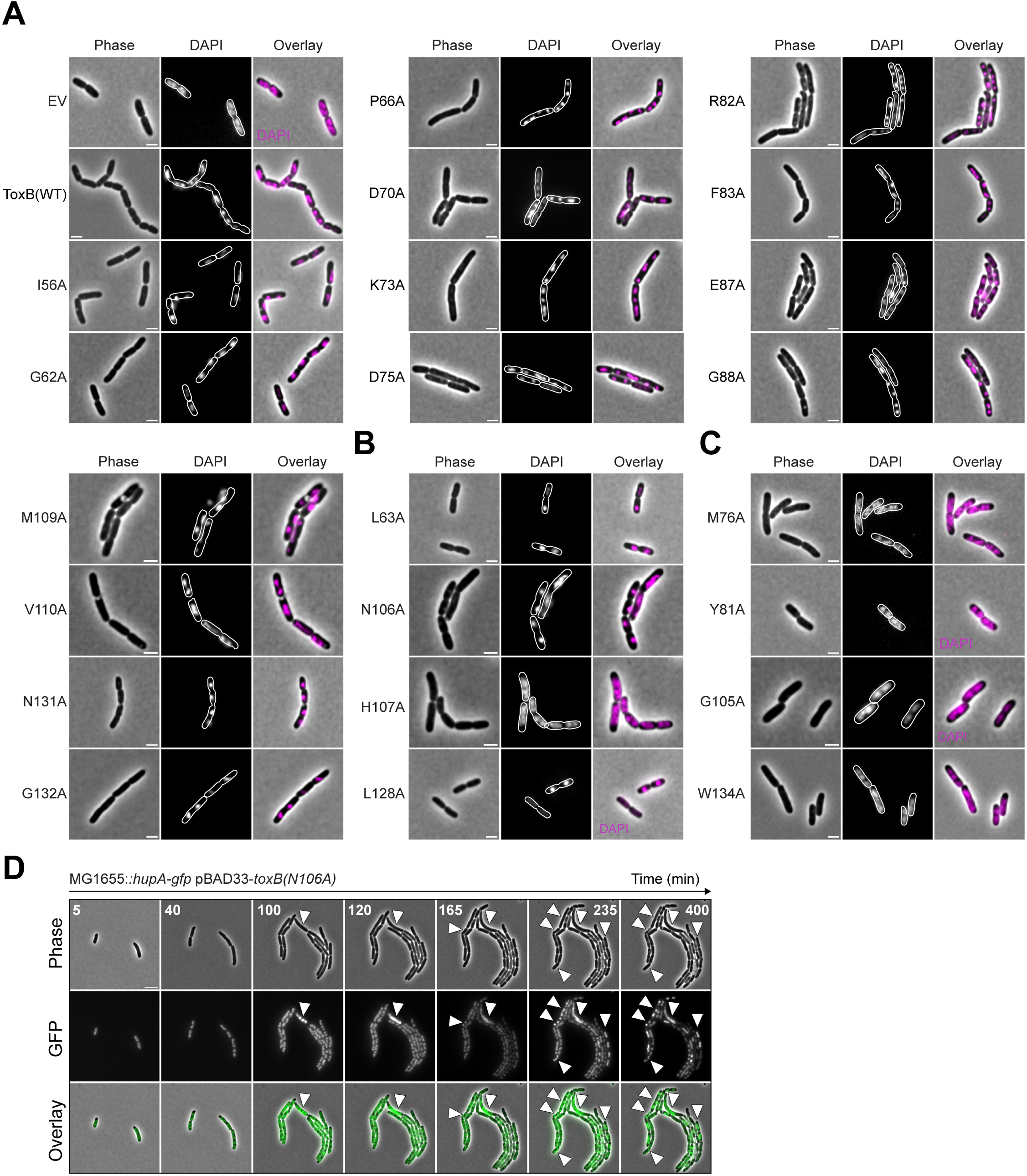
Single-cell analysis of nucleoid organization induced by ToxB alanine substitution mutants. Related to. **Figure 7. (A)** Toxic ToxB mutants induce nucleoid compaction similar to wild type. Representative phase-contrast, DAPI fluorescence, and overlay images of AB1157 *E. coli* cells producing ToxB(WT) or Class I alanine substitution mutants under inducing conditions. **(B)** Same as (A), only for Class II mutants (L63A, N106A, H107A, L128A), still showing nucleoid condensation comparable to wild-type ToxB. **(C)** Same as A-B, only for class III (non-toxic) mutants (M76A, Y81A, G105A, W134A), showing absence of nucleoid compaction. For all, cell outlines were generated by segmentation of phase-contrast images using Oufti. Scale bar is 2 µm. **(D)** A representative Class II mutant induces progressive nucleoid disorganization. Time-lapse microfluidics imaging of MG1655 cells expressing ToxB(H107A). Phase-contrast, HupA-GFP, and overlay images are shown at the indicated times. Arrowheads indicate progressive nucleoid disorganization during cell growth. Scale bar is 2 µm.

**Figure S7.**
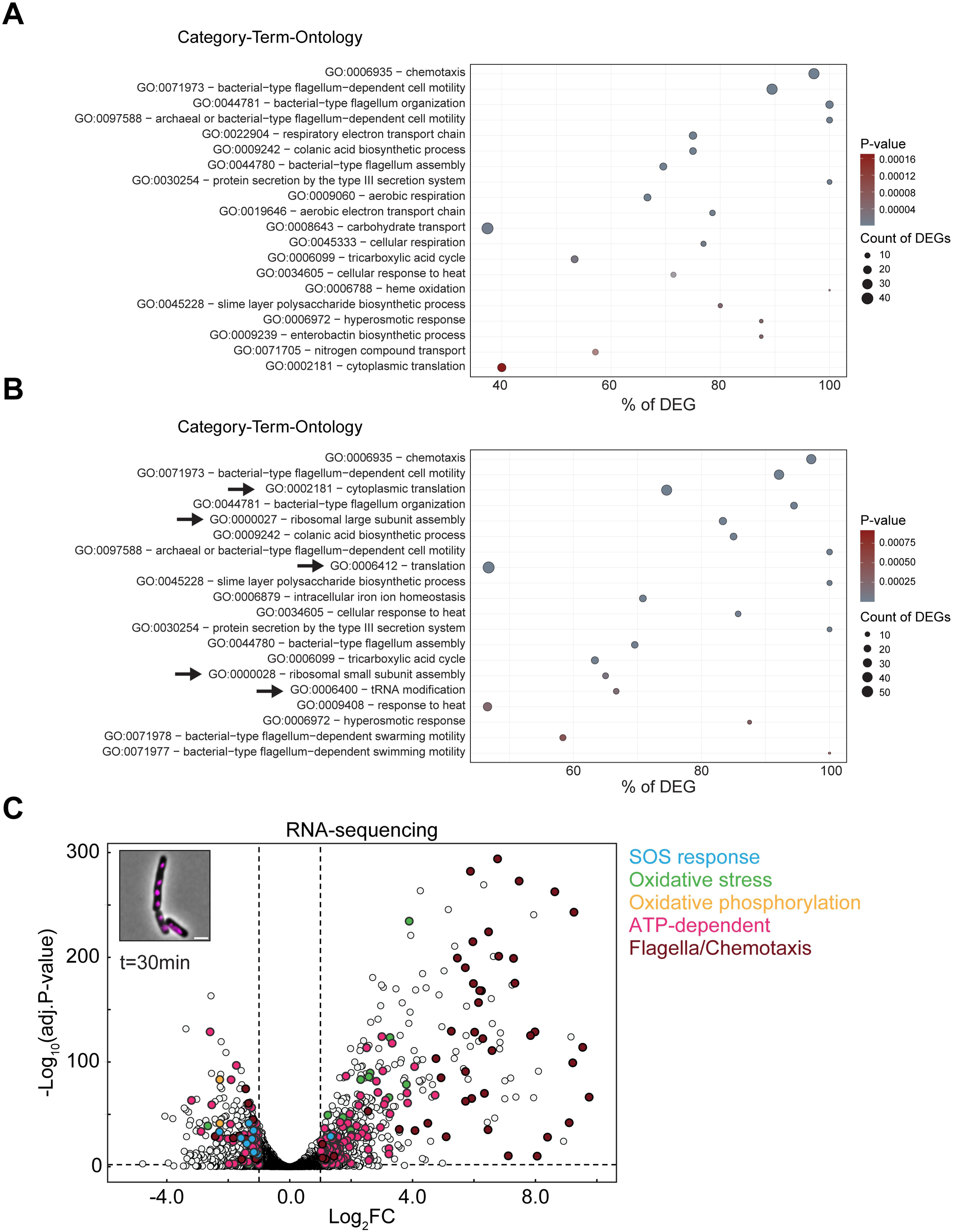
Reporter analyses and RNA-sequencing reveal early metabolic and envelope responses to ToxB intoxication. Related to. **Figure 8. (A)** Gene ontology (GO) enrichment analysis of differentially expressed genes (DEGs) identified by RNA-seq 15 minutes after induction of *toxB* (WT) in *E. coli* AB1157 compared with empty vector controls. Bubble plots show enriched GO biological process categories. The x-axis indicates the percentage of DEGs associated with each GO term, bubble size represents the number of genes assigned to the category, and bubble color reflects the adjusted *P* value. **(B)** GO enrichment analysis of DEGs identified 30 min after *toxB* (WT) induction. Bubble plot representation is as in panel A. Selected categories highlighted with arrows correspond to processes discussed in the main text. **(C)** RNA-seq volcano plot showing differential gene expression 30 min after induction of toxB (WT) compared with empty vector controls. The x-axis shows log₂ fold change (Log₂FC) and the y-axis shows −Log₁₀ adjusted *P* values. Genes associated with the SOS response (blue), oxidative stress (green), oxidative phosphorylation (orange), ATP-dependent processes (pink), and flagellar/chemotaxis functions (red) are highlighted. See also Supplementary Tables 2-5.

**Figure S8.**
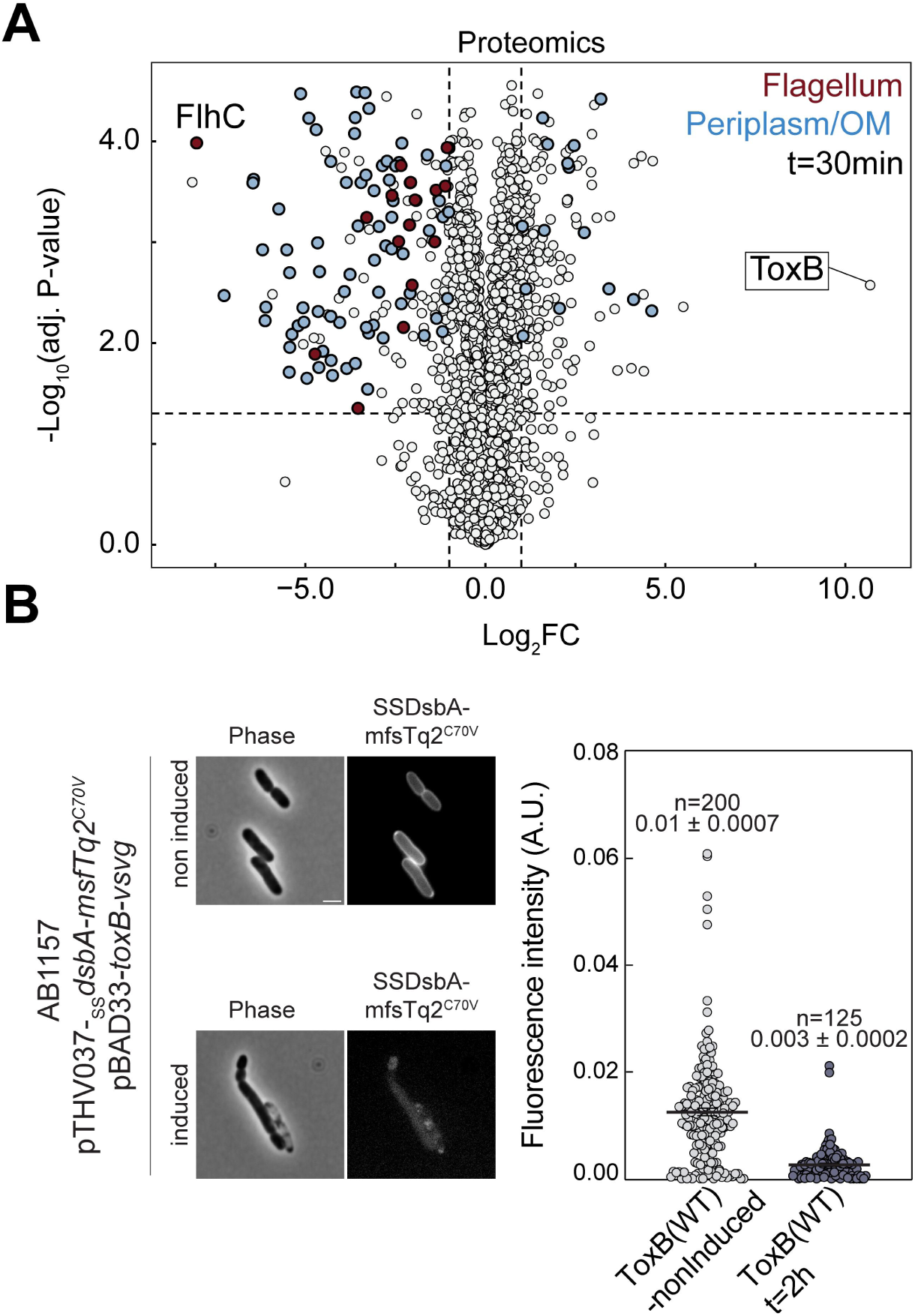
Proteomics reveal late envelope responses to ToxB intoxication. Related to. **Figure 8. (A)** Proteomics volcano plot showing differential protein abundance in *E. coli* AB1157 cells expressing ToxB (WT)compared with empty vector controls 30 minutes after induction. The x-axis shows log₂ fold change (log₂FC), and the y-axis shows −Log₁₀ adjusted *P* values. Proteins predicted to localize to the periplasm or outer membrane are highlighted in blue, while flagellar proteins are highlighted in red. See also Supplementary Tables 4-5. **(B)** Periplasmic reporter assay monitoring periplasm integrity during ToxB intoxication. Representative phase-contrast and fluorescence images of *E. coli* AB1157 cells expressing the DsbA signal peptide-*mfsTq2^C70V^*periplasmic reporter from plasmid pTHV037-*_SS_dsbA-mfsTq2^C70V^* together with pBAD33-*toxB-vsvg*. Cells were imaged under non-induced and induced conditions. Loss of periplasmic fluorescence signal is observed following toxin induction. Right: quantification of single-cell fluorescence intensity. Each point represents one cell. Mean fluorescence values ± stdv are indicated (n = 200 cells for non-induced and n = 125 cells for induced conditions).

